# Extracellular matrix protein composition dynamically changes during murine forelimb development

**DOI:** 10.1101/2020.06.17.158204

**Authors:** Kathryn R. Jacobson, Aya M. Saleh, Sarah N. Lipp, Alexander R. Ocken, Tamara L. Kinzer-Ursem, Sarah Calve

## Abstract

The extracellular matrix (ECM) is an integral part of multicellular organisms, connecting different cell layers and tissue types. During morphogenesis and growth, tissues undergo substantial reorganization involving cellular proliferation, migration, and differentiation. While it is intuitive that the ECM remodels in concert, little is known regarding how matrix composition and organization change during development. We utilized tissue fractionation and mass spectrometry to define ECM protein (matrisome) dynamics during murine forelimb development and resolved significant differences in ECM composition as a function of development, disease and tissue type. Additionally, we used bioorthogonal non-canonical amino acid tagging (BONCAT) to label newly synthesized ECM within the developing forelimb. We demonstrate the feasibility of using BONCAT to enrich for newly synthesized matrisome components and identified differences in ECM synthesis between morphogenesis and growth. This resource will guide future research investigating the role of the matrisome during complex tissue development.

## Introduction

After fertilization, the cells of the embryo transition from a homogenous cell mass to a heterogeneous collection of tissues that assemble into the organ systems of the adult. Techniques characterizing the genome, epigenome, transcriptome and proteome dynamics during embryonic development have provided a global understanding of signaling pathway dynamics that regulate cellular function during morphogenesis (Yang, et al., 2019; Zheng and Xie, 2019; Baralle and Giudice, 2017; Scialdone, et al., 2016). Another important regulator of embryogenesis is the extracellular matrix (ECM), a collection of proteins and glycosaminoglycans that assemble into tissue-specific, interpenetrating networks. These active networks maintain the physical integrity of tissues, serve as a reservoir for growth factors and act as a medium for sensing and transducing mechanical signals through ECM-cell interactions (Felsenthal and Zelzer, 2017; Rozario and Desimone, 2010). As the demands of the environment change during morphogenesis, growth and homeostasis, it is likely the ECM remodels in concert; however, the contribution of the ECM, and how it changes during complex tissue development, is not well defined.

The murine forelimb is an ideal model system to investigate how different factors regulate morphogenesis. In the developing mouse, the forelimb bud starts off as a relatively disorganized tissue and, over the course of 4-5 days, the cells assemble into muscle, connective tissue and cartilage, with seamless, complex interfaces to form the basis of a functional musculoskeletal system (Huang, 2017; Charvet, et al., 2012). The ECM composition and architecture varies widely across these distinct tissues; however, the influence of the matrix during the specification, integration and maturation of musculoskeletal tissues is poorly understood.

The gap in knowledge regarding ECM dynamics during morphogenesis can be attributed to the previous lack of techniques that could resolve changes in the matrix within developing tissues. The identification of differences in ECM composition between tissues was hindered by the relative insolubility of the matrisome compared to intracellular proteins (Naba, et al., 2012). Additionally, ECM proteins make up a small percentage of the total protein content in many tissues. To address these challenges, techniques combining liquid chromatography-tandem mass spectrometry (LC-MS/MS) and tissue fractionation were developed to identify how the composition of ECM proteins, or the matrisome, varies between different adult and pathological tissues (reviewed in (Taha and Naba, 2019)). Recently, our lab extended these techniques to analyze the matrisome of embryonic murine tissues (Saleh, et al., 2019a); however, this only provided a snapshot of the static matrisome at a certain stage of development and was unable to resolve when specific proteins were made. Identification of the proteins synthesized at a given timepoint will provide more information about the dynamics of critical components that drive changes in tissue structure and remodeling during development.

Biorthogonal non-canonical amino acid tagging (BONCAT) is a technique previously used by our group and others to identify the newly synthesized proteome in embryonic, adult and pathological tissues (Saleh, et al., 2019b; Dieterich, et al., 2006). BONCAT utilizes non-canonical amino acids (ncAAs) that are incorporated into proteins using the endogenous cellular translational machinery. These ncAAs possess biorthogonal moieties (*e.g.* azides) that enable the enrichment of newly synthesized proteins (NSPs) through click chemistry reactions with complementary chemical groups (*e.g.* alkynes). BONCAT was previously used to enrich for soluble proteins, but translation of this technique to the insoluble matrisome has yet to be demonstrated.

The goals of this study were to (1) expand our recent work on the proteomic analysis of developing tissues to account for significant increases in protein content during embryogenesis; (2) establish a baseline proteomic resource of static and newly synthesized matrisome dynamics during morphogenesis and growth in the wild-type (WT) murine forelimb; and (3) resolve differences in matrisome composition between (a) pathological and WT tissues and (b) tissues from distinct organ systems. We were able to resolve differences in relative matrisome composition of forelimbs that ranged from embryonic day (E)11.5-postnatal day (P)35, and found that ECM protein composition during morphogenesis (E11.5-E14.5) was markedly different compared to growth timepoints (P3-P35). To identify NSPs, mice were tagged with the ncAA azidohomoalanine (Aha) when musculoskeletal patterning was just established (E13.5-E14.5) and compared to an adolescent timepoint representative of growth (P35 – P36). The standard BONCAT method was optimized to enrich for Aha-tagged ECM prior to LC-MS/MS and identify different patterns of synthesis that could not be revealed by only investigating the static matrisome. We next confirmed that the tissue fractionation and LC-MS/MS techniques were able to resolve differences in matrisome composition in WT E14.5 murine forelimbs compared to *Col1a2*-mutants, which exhibit musculoskeletal defects. As expected, we found a significant decrease in the abundance of chains associated with type I collagen fibrils (COL1A1, COL1A2, COL3A1 and COL5A1). Interestingly, there was an increase in lysyl oxidase-like (LOXL) proteins that modify constituents of collagen fibrils, which can lead to over-modification and incorrect integration of these chains into fibrils. Finally, these techniques were used to determine the unique composition of the matrisome in WT E14.5 brains compared with forelimbs. While the most abundant ECM proteins found in both tissues were similar, there were many exclusively identified in each tissue, which correlated with the contrasting biological functions. Collectively, these results demonstrate the feasibility of using tissue fractionation and ncAA tagging to resolve differences in ECM protein composition as a function of age, disease and tissue type, providing a map of the matrisome during forelimb development. These methods can be easily extended to investigating the role of the matrisome in other systems during morphogenesis and growth.

## Results and Discussion

### Fractionation of whole embryonic tissue increased ECM protein identification and enabled temporal analysis of the matrisome

We previously demonstrated that fractionation of E15.5 murine embryos facilitated the analysis of the matrisome via LC-MS/MS (Saleh, et al., 2019a). To confirm that fractionation provides the same benefits for earlier embryonic timepoints wherein the ECM may be less cross-linked, E12.5 wild-type (WT) embryos were homogenized then fractionated using buffers designed to selectively extract cytosolic (C), nuclear (N), membrane (M) and cytoskeletal (CS) proteins, leaving behind an ECM-rich insoluble (IN) pellet (Naba, et al., 2015) (**Figure S1A)**. All samples were processed for LC-MS/MS, raw protein intensities were determined by MaxQuant (Cox and Mann, 2008) and proteins were categorized into cellular compartments (Saleh, et al., 2019a). Within the homogenate, 20 ECM proteins were identified; however, these proteins only represented 0.8% ± 0.3% of the total raw intensity (**Figure S1B, Table S1**). In contrast, the percentage of matrisome in the IN pellet was 91.5% ± 3.5%, indicating tissue fractionation significantly increased the identification of ECM proteins in E12.5 embryos (**Table S1**). The amount of ECM protein in the C, N and M fractions contributed to less than 1.0% ± 0.9% of the total raw intensity. In comparison, ECM proteins contributed 2.3% ± 1.1% of the total raw intensity of the CS fractions. For subsequent studies, the CS and IN fractions were analyzed via LC-MS/MS separately, while the C, N and M fractions were combined into one ‘CNM’ fraction.

To investigate whether fractionation was amenable for temporal analysis of the embryonic matrisome, the CNM, CS and IN fractions of E11.5-E14.5 WT embryos were analyzed by LC-MS/MS (**Figure 1A**). Across all fractions and timepoints, 3227 proteins were identified, 122 of which were part of the matrisome (**Table S2**). Less than 1% and 5% of the raw intensity in CNM and CS fractions, respectively, were attributed to ECM proteins, consistent with our previous study (Saleh, et al., 2019a) (**Figures 1B, S1**). Importantly, the matrisome was enriched in the IN fraction, where at least 30% of the raw intensity was from ECM proteins.

**Figure 1.**
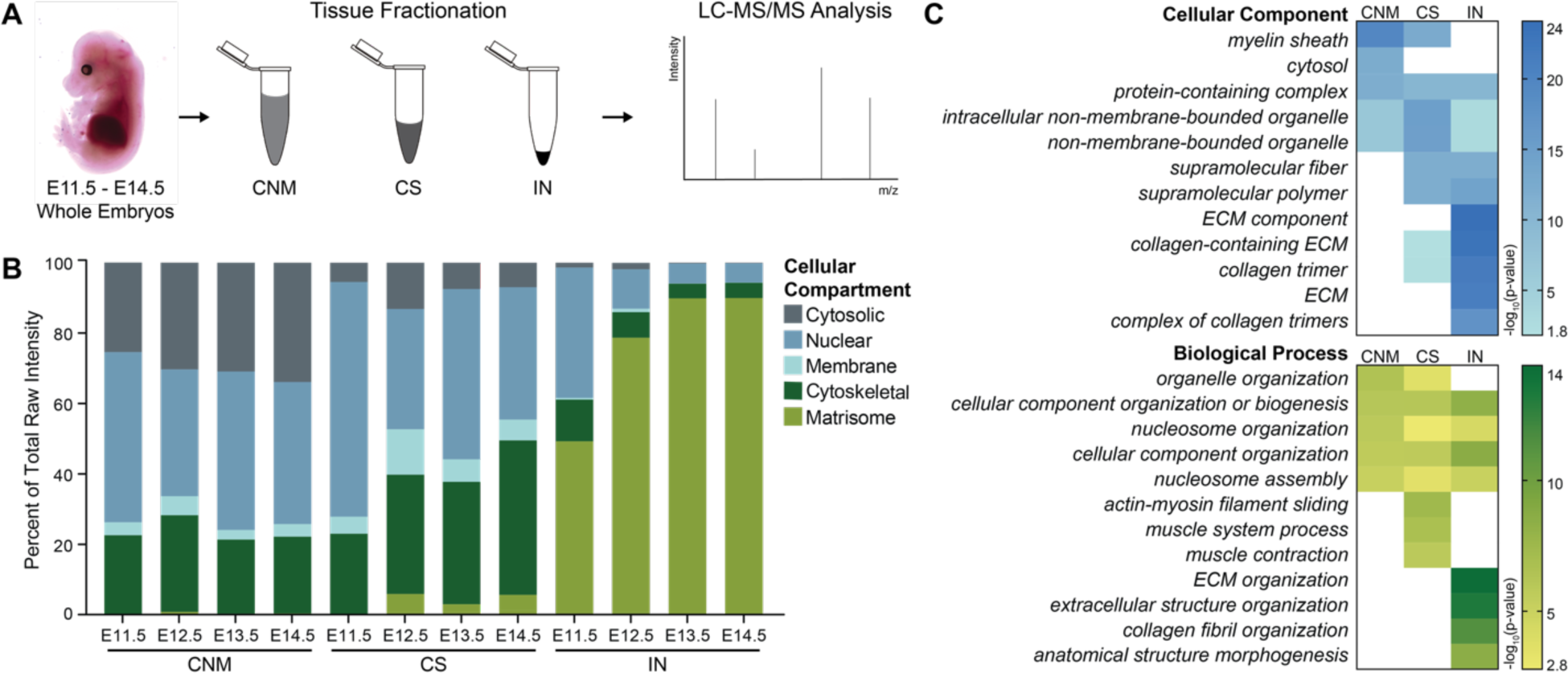
Proteomic analysis of whole murine embryos. **(A)** Tissue fractionation was combined with LC-MS/MS to analyze E11.5-E14.5 whole embryos (*n*=3 biological replicates/timepoint). Cytosolic (C), nuclear (N) and membrane (M) fractions were combined into one CNM fraction. CNM, CS and IN fractions were analyzed by LC-MS/MS and raw intensities were determined using MaxQuant. **(B)** The distribution of cellular compartments, as defined by (Saleh, et al., 2019a), in CNM, CS and IN fractions plotted as average across biological replicates. Two-way ANOVA revealed the percentage of total raw intensities attributed to the matrisome was significantly different between timepoints (*p*<0.0001) and fractions (*p*<0.0001). **(C)** The top 5 significant GO “Cellular Component” and “Biological Process” terms generated from the 50 most abundant proteins within each CNM, CS and IN fraction of E14.5 embryos indicated successful enrichment of ECM-related GO terms in the IN fraction (see also **Figure S2**).

GO analysis was conducted on the 50 most abundant proteins in each fraction for E14.5 embryos. “Cellular Component” terms indicated proteins extracted in the C, N and M buffers were from intracellular compartments, and the most significant “Biological Process” terms corresponded to intracellular organization and protein interactions (**Figure 1C**). In contrast, GO terms generated by the most abundant proteins of the IN fraction highly correlated with the ECM. Unbiased hierarchal clustering for the same set of proteins showed matrisome components were enriched in the IN fraction (**Figure S2**). Collectively, these results validate that fractionation enhanced identification of the matrisome in developing murine embryos.

### Global ECM composition significantly changed during forelimb development

To investigate matrisome dynamics as a function of musculoskeletal development, entire forelimbs, from the scapula through the digit tips, were isolated from E11.5-P35 mice (**Figure 2A**). Tissues were fractionated as described above and CS and IN fractions were analyzed by LC-MS/MS (**Figure 2B**). Collectively, 1601 proteins were identified, including 147 ECM proteins (**Table S3**). The amount of matrisome varied significantly between timepoints and fractions; similar to whole embryonic tissue, ECM proteins were significantly enriched in the IN fraction (**Figure 2C**).

**Figure 2.**
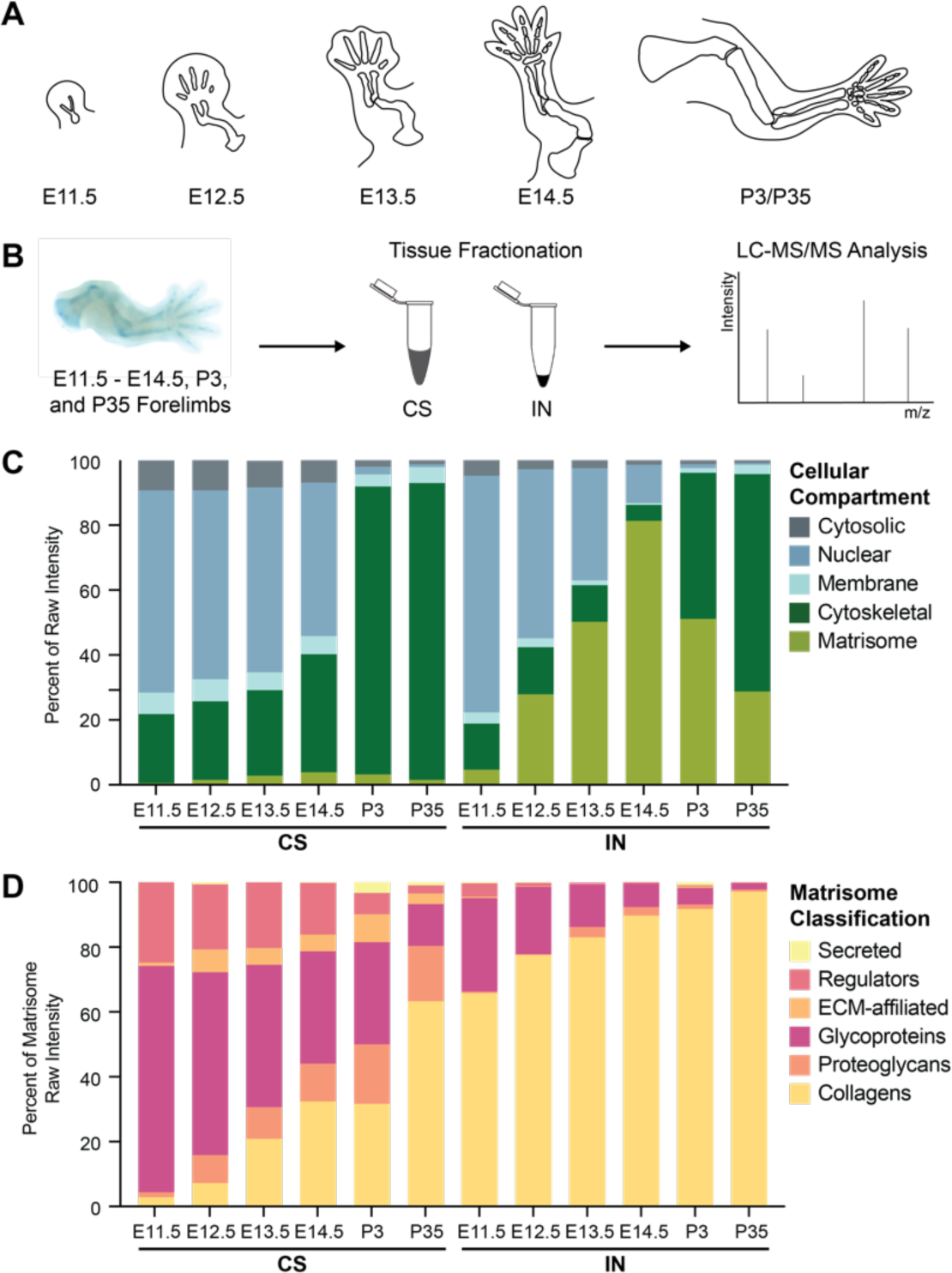
Proteomic analysis of developing murine forelimbs. **(A)** Architecture of forelimbs dissected from embryos and pups (not drawn to scale). **(B)** Tissue fractionation was combined with LC-MS/MS analysis to investigate matrisome content of CS and IN of E11.5-P35 forelimbs (*n*=3 biological replicates). **(C)** The distribution of cellular compartments, plotted as an average percentage of raw intensity, identified in CS and IN fractions. Two-way ANOVA showed the percentage of matrisome was dependent on timepoint (*p*<0.0001) and fraction (*p*<0.0001). **(D)** ECM proteins were categorized as defined by (Naba, et al., 2012), and percentages of raw matrisome intensity were plotted as a function of development. Three-way ANOVA revealed the distribution of matrisome components was significantly influenced by timepoint and fraction (*p*<0.0001).

Collagens were the most abundant ECM proteins identified in the IN fraction at all timepoints (**Figure 2D**). ECM glycoproteins were the second most abundant, but the relative percentage of the total matrisome decreased with development. Similar trends were found in the CS fractions; however, there were significantly more secreted factors, ECM regulators and ECM-affiliated proteins compared to the IN fraction (**Table S3**).

### Matrisome dynamics varied during musculoskeletal morphogenesis and growth

Next, we combined protein intensities from CS and IN fractions and analyzed matrisome changes during morphogenesis and growth using a heat map (**Figure 3A**). Combining LFQ intensities from CS and IN fractions allowed for direct comparison between timepoints (**Figure S3C**). The amount of matrisome extracted in each fraction varied depending on ECM classification (**Figure S3D, E**). Fibril-forming collagens (types I, II, III, V, XI) and ECM glycoproteins, in particular those proteins that make up elastic fibrils (ELN, FBLN2, FBN1, FBN2, MFAP2), were predominantly found in the IN fraction, which is likely due to the extensive cross-linking that holds these proteins together (Schräder, et al., 2018; Tanzer, 1973). Interestingly, many proteoglycans, including small leucine-rich proteoglycans (SLRPs), were found at larger percentages in the CS fraction. SLRPs are known to play roles in regulating collagen fibril assembly and bind to fibrils non-covalently, which may explain the increased extractability of these proteins (Neame, et al., 2000). The majority of the ECM-affiliated proteins, ECM-regulators and secreted factors were identified either exclusively or more abundant in the CS fraction. Many of these proteins remain in cytoplasmic regions until needed for matrix-cell signaling transduction (MEGF6), collagen post-translational modifications (LOX, PLOD1-3) and network degradation (MMP13) (Vallet and Ricard-Blum, 2019; Hu, et al., 2018; Qi and Xu, 2018; Lu, et al., 2011). Since these proteins enzymatically modulate the existing ECM, rather than integrate into the networks, they were are more soluble and removed in earlier fractions. While many of the recent matrisome studies only analyze the final, IN fraction (Fava, et al., 2018; Suna, et al., 2018; Naba, et al., 2017a), our data showed that some ECM proteins are extracted in the other buffers. Depending on the proteins of interest, future investigations need to ensure the appropriate fractions are analyzed. Nevertheless, fractionation enables identification of how ECM changes, here as a function of age (**Figure 3**), even if only one fraction is analyzed.

**Figure 3.**
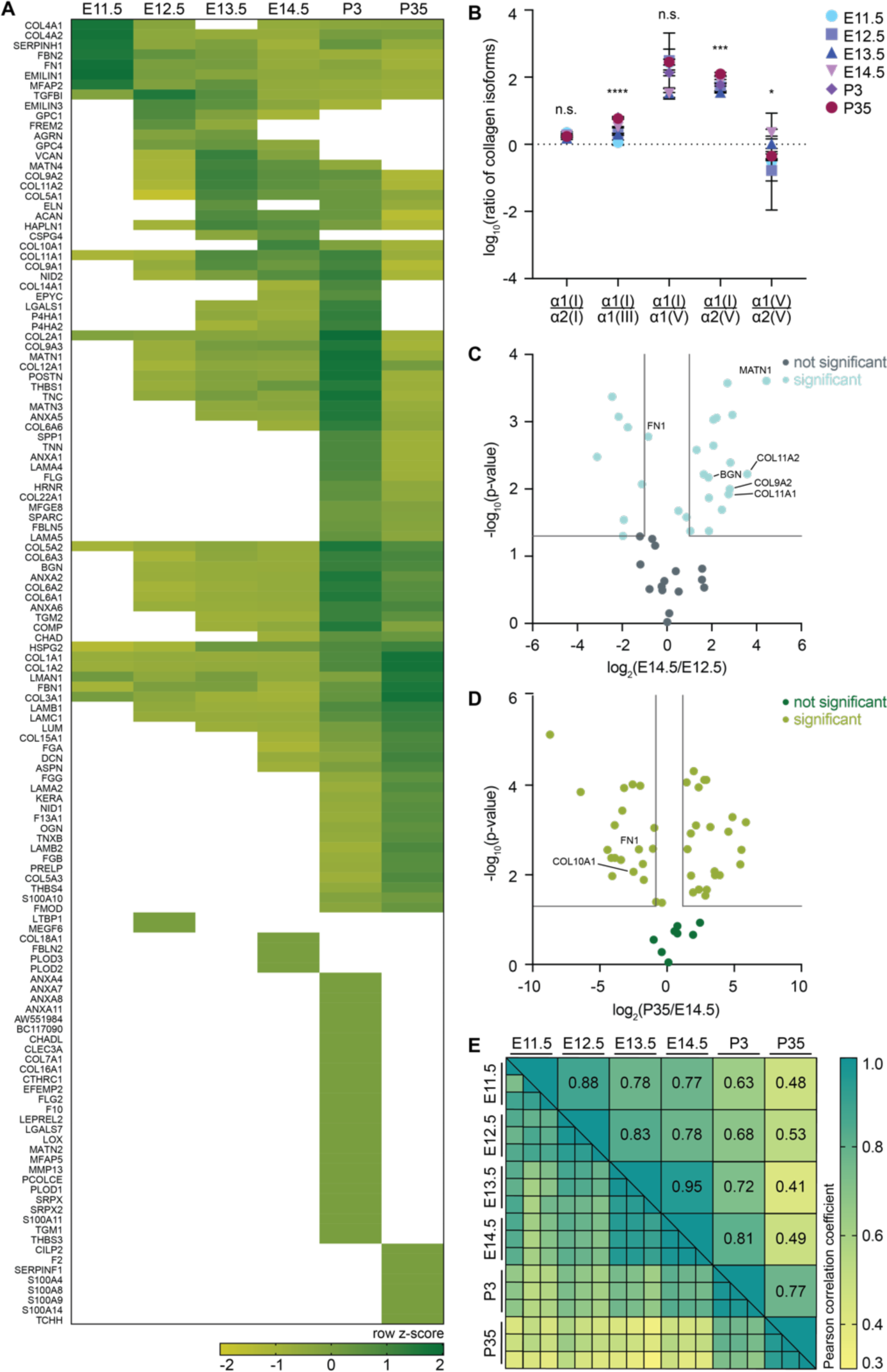
ECM protein composition varies as a function of murine musculoskeletal development. E11.5-P35 WT forelimbs were analyzed using LC-MS/MS as described in **Figure 2**. (**A**) LFQ intensities were normalized and combined from both CS and IN fractions for each timepoint (see **Methods**) and averaged across biological replicates (*n*=3). Proteins were clustered, based on row z-score, to show dynamics as a function of development. Proteins identified in *n*≥2 biological replicates were included in the heat map analysis. White boxes signify zero intensity values. **(B)** The ratios of the raw intensities of collagen chains involved in type I collagen fibrillogenesis significantly varied between timepoints. Statistical differences were determined for each ratio by one-way ANOVA across timepoints (n.s. denotes *p*>0.05; \**p*<0.05; \*\*\**p*<0.001; \*\*\*\**p*<0.0001) (**Table S3**). **(C, D)** Volcano plots comparing normalized LFQ intensity values of ECM proteins identified at E12.5 and E14.5 or E14.5 and P35. Grey lines denote ≥2-fold change and *p*<0.05 (two-tailed t-test). **(E)** Pearson correlation coefficients comparing the matrisome between timepoints.

In the early stages of morphogenesis, key proteins are critical for various ECM networks to polymerize and facilitate the formation of distinct tissues. ECM proteins associated with morphogenesis (FN1, FBN2, EMILIN1, TGFβI) were abundant in the forelimb and expression decreased at older timepoints (Schiavinato, et al., 2017; Xu, et al., 2009; Sottile and Hocking, 2002; Chaudhry, et al., 2001) (**Figure 3A, Table S3**). For example, the normalized LFQ intensity of FN1 was high at E11.5 then decreased in relative abundance as a function of forelimb development (**Figure 3)**. FN1 orchestrates the assembly of ECM protein networks, including types I and III collagen, fibrillins, THBS1 and LTBP1 (Kadler, et al., 2008; Dallas, et al., 2005), and global FN1 knockout is embryonic lethal before E11 (George, et al., 1993).

Type I collagen fibrils are another important component of tissues. The relative amount of α1 and α2 chains that comprise the type I collagen heterotrimer (α1(I) and α2(I), respectively), significantly increased with development and accounted for 42.0% ± 4.1% – 84.8% ± 2.1% of total collagen content (**Table S3**). The ratio between α1(I) and α2(I) chains was not significantly different between timepoints and the average across all timepoints (1.8 ± 0.3) was similar to the standard ratio of 2:1 for α1(I):α2(I) reported in the literature (Birk and Brückner, 2011) (**Figure 3B, Table S3**).

In contrast, the relative abundance of collagens involved in type I collagen fibrillogenesis, including types III and V (Kadler, et al., 2008), significantly varied over development. Type III collagen forms independent fibrils out of α1(III) homotrimers as well as heterotypic fibrils with type I collagen (Liu, et al., 1997). *In vivo* and *in vitro* studies suggest that type III collagen inhibits the increase in diameter of type I collagen fibrils (Asgari, et al., 2017; Liu, et al., 1997). The ratio of α1(I):α1(III) significantly increased between E11.5 and P35 (**Figure 3B, Table S3**), consistent with previous observations that type I collagen fibril diameter increases during development (Parry, et al., 1978).

Type V collagen is a quantitatively minor component, accounting for 1.1% ± 0.5% of collagen content across all timepoints (**Table S3**). Nevertheless, it is critical for the initiation of type I collagen fibrillogenesis and forms homo- and heterotypic combinations of three alpha chains, α1(V), α2(V) and α3(V) (Wenstrup, et al., 2004). The ratio of α1(I):α1(V) did not significantly change as a function of development; however, α1(I):α2(V) and α1(V):α2(V) were significantly different between stages of morphogenesis and growth (**Figure 3B, Table S3**). Variations in the chain composition of type V collagen were previously described in the cornea (Birk, 2001); however, additional studies need to be conducted to elucidate the functional importance of changes in type V collagen isoforms during forelimb development.

Other distinct trends were visualized by comparing protein intensities at two timepoints during morphogenesis (E12.5 vs. E14.5) and morphogenesis and growth (E14.5 vs. P35) using volcano plots. ECM proteins associated with chondrogenesis were increased in E14.5 forelimbs compared to E12.5 (**Figure 3C**), including types IX and XI collagen, MATN1 and BGN (Li, et al., 2018; Krug, et al., 2013; Embree, et al., 2010; Nicolae, et al., 2007). Proteins important during endochondral ossification, such as type X collagen (Kwan, et al., 1997), were significantly more abundant at E14.5 and decreased at P35 (**Figure 3D**), indicating that the transition from cartilage to bone neared completion by P35. Furthermore, ECM proteins identified exclusively, or in higher abundance, at P35, such as CILP2 and OGN, are associated with bone maturation and homeostasis (Lee, et al., 2018; Bernardo, et al., 2011).

Comparison of Pearson correlation coefficients confirmed the difference in matrisome content between embryonic and adolescent timepoints (**Figure 3E**). There was low correlation between embryonic (E11.5-E14.5) and P35 forelimbs, whereas the correlation within embryonic timepoints was higher. Taken together, these data demonstrated ECM proteins are differentially expressed during musculoskeletal tissue morphogenesis and growth.

### Spatial distribution of ECM proteins changed within musculoskeletal tissues during development

LC-MS/MS analysis revealed how the ECM dynamically changes in the developing forelimb (**Figure 3**); however, proteomics alone does not provide spatial information of the matrisome within distinct tissues. To gain more insight into how the distribution of the matrisome changes during forelimb development, and validate the LC-MS/MS results, ECM proteins with differing patterns of abundance were investigated via immunohistochemistry (IHC, **Figure 4, Table S4**). While the distribution of the mRNA or protein for some of these ECM can be found in the literature at distinct stages of development, a comprehensive investigation of the matrisome during musculoskeletal tissue assembly during the timepoints in this study has not been conducted. The identification of these ECM proteins with respect to specific tissues is summarized in **Table S4**.

**Figure 4.**
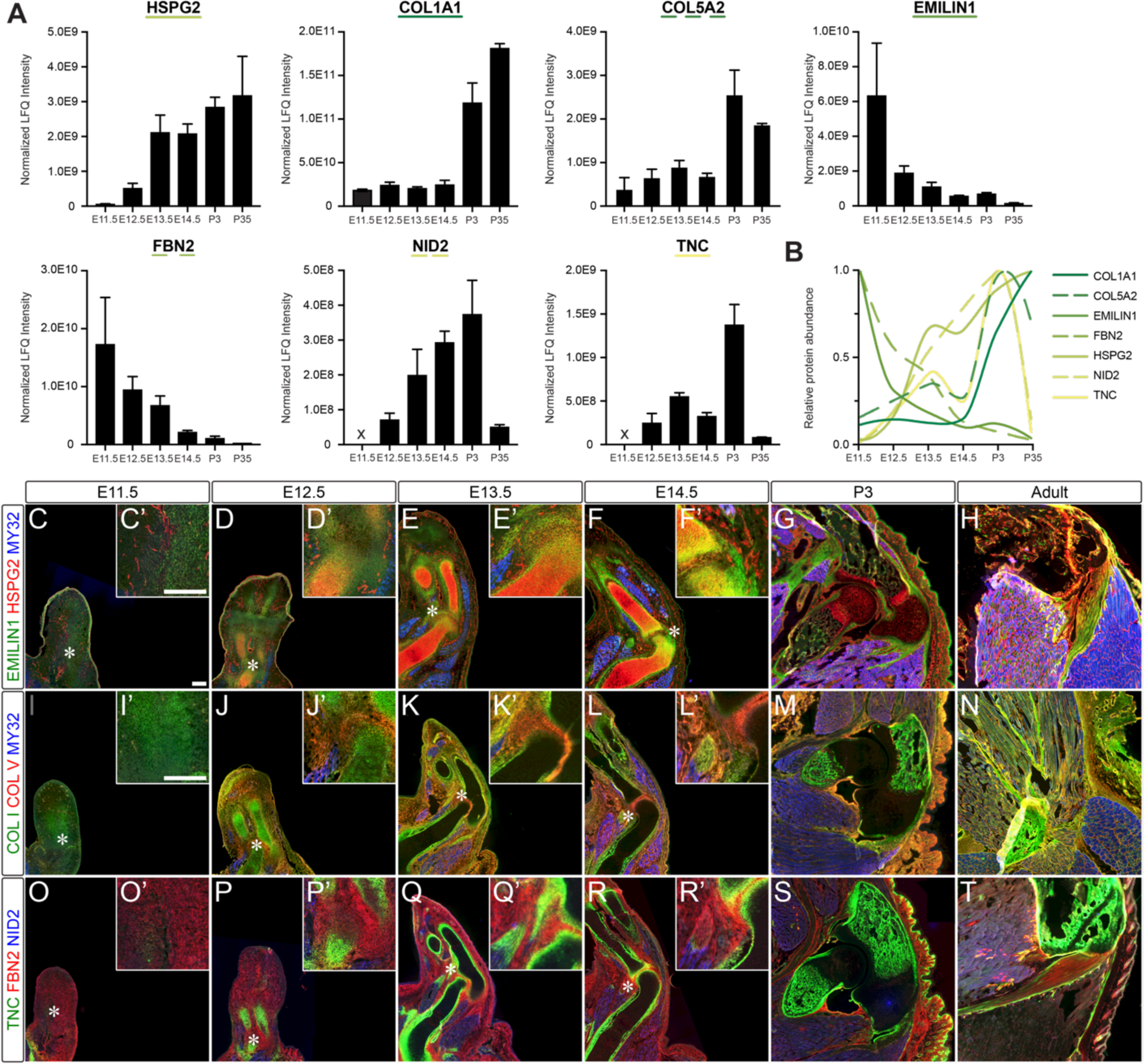
The ECM is differentially distributed within musculoskeletal tissues during forelimb development. **(A)** Normalized combined LFQ intensities from WT E11.5-P35 forelimbs for proteins selected for immunohistochemistry plotted as average (*n*=3 biological replicates). Intensity values and one-way ANOVA statistics for each protein are reported in **Tables S3. (B)** Graphical summary of protein dynamics displayed in (**A**). The largest value for each protein was set to 1 and the remaining were scaled to show relative abundance as a function of timepoint. **(C-T)** Cryosections from E11.5-E14.5, P3 and adult forelimbs were stained with antibodies against: **(C-H, C’-F’)** EMILIN1 (green), HSPG2 (red), and myosin heavy chain, a marker for differentiated skeletal muscle (MY32; blue); **(I-N, I’-L’)** type I collagen (COL I; green), type V collagen (COL V; red), and MY32 (blue); **(O-T, O’-R’)** TNC (green), FBN2 (red), and NID2 (blue). Insets (indicated with’) are a 3× enlargement of the region containing the nascent elbow (*) for E11.5-E14.5. Scale bars=200 µm. Individual channels and secondary antibody only negative control panels are shown in **Figures S4-S6**.

HSPG2 and type I collagen (COL I) were two proteins that continuously increased in abundance between E11.5 – P35 (**Figure 4A, Table S4**). Staining for HSPG2 localized to blood vessels at E11.5, but was abundant in the skeletal template in all subsequent timepoints, with more diffuse labeling throughout the surrounding mesenchyme (**Figures 4, S4**). Postnatally, HSPG2 increased throughout all mesenchymal tissues. HSPG2 is a component of the pericellular matrix surrounding mature muscle fibers, tenocytes, chondrocytes and osteocytes (Martinez, et al., 2018; Taylor, et al., 2011; Hassell, et al., 2002). The importance of this proteoglycan in the musculoskeletal system is seen in Schwartz-Jampel Syndrome, which is due to the knockdown of HSPG2, and manifests in chondrodysplasia and neuromyotonia (Stum, et al., 2008). In contrast, COL I was diffusely distributed throughout the limb bud, with slightly increased expression in condensing cartilage at E11.5-E12.5 (**Figures 4, S5**). However, at E13.5 there was little COL I visualized in the skeletal template compared with the surrounding mesenchyme until the onset of endochondral ossification. COL I was widely expressed in the adult, as expected since COL I is the most abundant ECM protein in mammals where mutations lead to defects, such as osteogenesis imperfecta, in which the mechanical integrity of tissues is compromised (Di Lullo, et al., 2002).

Type V collagen (COL V), which facilitates COL I fibrillogenesis, also increased in abundance as a function of development (**Figure 4A**). The distribution of COL V was similar to that of COL I, except it was not localized to the condensing cartilage at E12.5. In addition, it was enriched at the cavitating elbow joint, whereas COL I was more prevalent in the rest of the perichondrium/osteum. This is consistent with a previous observation that *pro-Col5a3* mRNA was highly expressed at the hip joint in E15.5 embryos (Imamura, et al., 2000). Notably, defects in COL V chains lead to Ehlers-Danlos Syndrome, which is characterized by laxity of connective tissues, particularly around the joint where the insertions of COL I- and COL V-rich tendons and ligaments form (Symoens, et al., 2012; Imamura, et al., 2000).

Two proteins that were abundant at E11.5 and decreased over development were EMILIN1 and FBN2 (**Figure 4A**). Similar to COL I, both were diffuse throughout the limb at E11.5; however, the distribution became more restricted to the perichondrium/osteum, tendons and ligaments over time (**Figures 4, S5, S6**). In the adult, EMILIN1 and FBN2 were found at lower levels, compared with earlier timepoints, with the exception of the tendons and ligaments (**Figure 4**). The abundance of FBN2 and EMILIN1 early in limb morphogenesis indicates these ECM proteins play important roles in musculoskeletal patterning. Indeed, knockout of FBN2 leads to abnormal mesenchymal differentiation and syndactyly (Arteaga-Solis, et al., 2001), and mutation of *EMILIN1* leads to musculoskeletal deformities (Capuano, et al., 2016).

NID2 and TNC are two proteins that transiently peak in abundance during development; both increased from E12.5-P3, but then significantly decreased between P3-P35 (**Figure 4A, Table S4**). NID2, a basement membrane protein, surrounded developing myotubes at E12.5 and then had a more punctate distribution at later timepoints (**Figure 4, S6**). This staining pattern is consistent with reports that NID2 becomes restricted to the neuromuscular junction during maturation (Fox, et al., 2008). TNC was first visualized in the condensing cartilage at E12.5 and then was predominantly found in the perichondrium/osteum, tendons, ligaments and bony spicules. While the restriction of TNC to the perichondrium/osteum is similar to that seen for FBN2 and EMILIN1, knockout of TNC does not lead to any overt musculoskeletal defects (Mackie and Tucker, 1999). Nevertheless, the transient upregulation of TNC during muscle repair and regeneration suggests this protein plays a currently undefined role in tissue growth that may be compensated for by other ECM at earlier timepoints (Calve, et al., 2010; Fluck, et al., 2008).

Overall, visualization of various ECM via IHC correlated with the changes in abundance identified with LC-MS/MS. Notably, ECM with similar changes in abundance were differentially localized within the limb, highlighting the need to not only identify what matrisome components are present but the distributions within tissues as well.

### Analysis of newly synthesized proteins at distinct timepoints identified dynamic changes in matrisome composition

While LC-MS/MS and IHC analyses identified matrisome components that are present at a given timepoint, the proteins that are actively being synthesized at a specific stage cannot be distinguished. We hypothesized that identification of NSPs using BONCAT would reveal differences in the distribution of NSPs during morphogenesis compared to growth timepoints. Pregnant females were injected at E13.5 with either Aha or PBS (control) and forelimbs were harvested from embryos 6- and 24-hours post injection (hpi; E13.75 and E14.5, respectively) to identify ECM proteins actively synthesized during morphogenesis. To identify matrisome synthesized during growth, non-pregnant female mice were injected with either Aha or PBS at P35 and harvested 6- and 24-hpi (P35.25 and P36, respectively). Forelimbs were fractionated and each IN fraction was split into two samples: (1) an “unenriched” sample that represented the background, static proteome, analogous to the IN fractions described in **Figure 2**, and (2) an “enriched” sample containing the isolated Aha-labeled proteins (**Figure 5A**). To enrich for and identify Aha-labeled ECM, it was necessary to resuspend the IN fraction in a strong denaturing agent (8M urea) to partially solubilize the relatively insoluble matrisome. The resuspended pellets were subjected to brief cycles of ultrasonic energy to further enhance the solubilization of ECM proteins. In addition, proteins were deglycosylated with chondroitinase ABC to digest the chondroitin sulfate chains and increase the accessibility of Aha residues to the biotin alkyne linker. This optimized technique enabled the use of BONCAT to characterize the *in vivo* matrisome for the first time.

**Figure 5.**
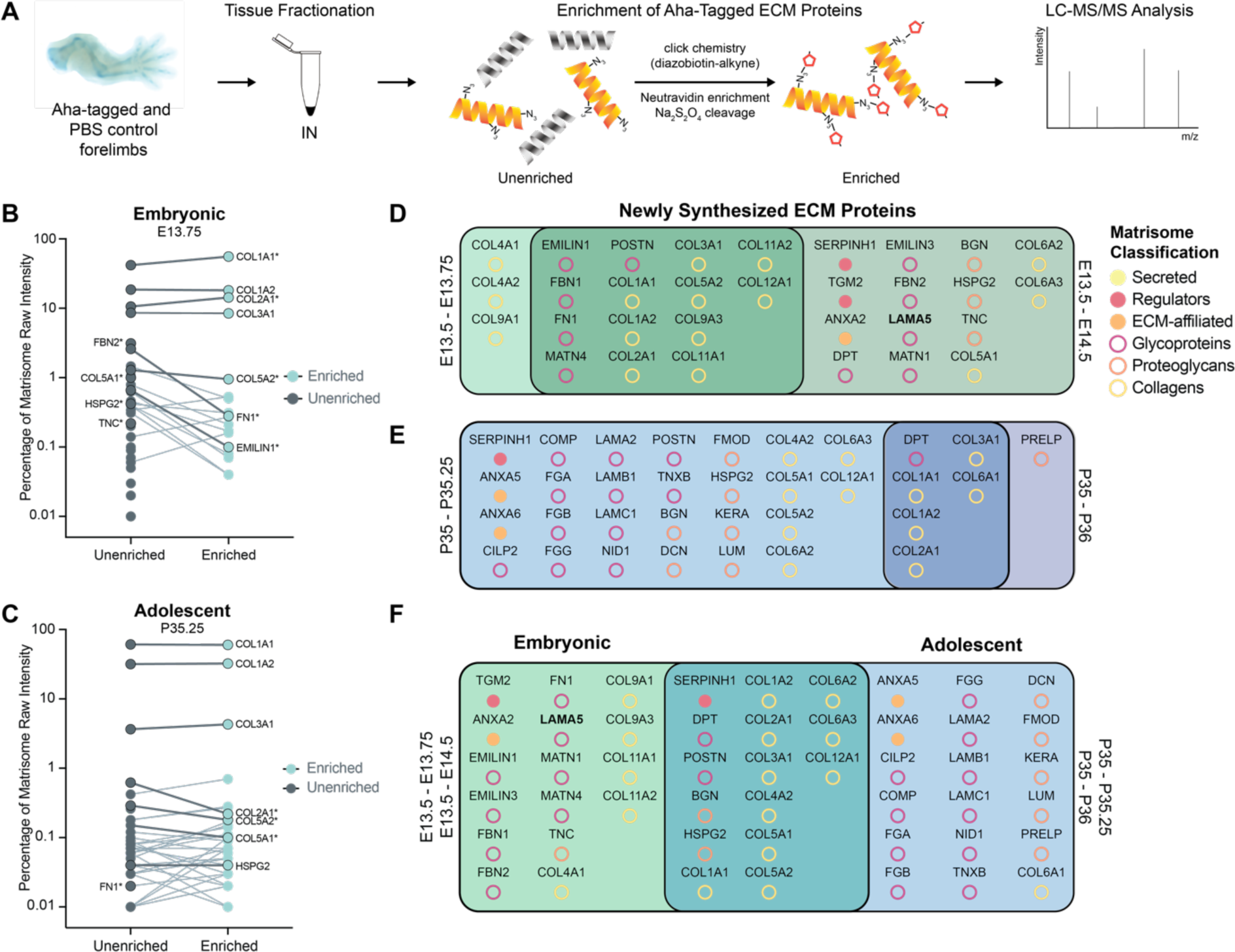
Newly synthesized ECM proteins vary during morphogenesis and growth. (**A**) *In vivo* Aha-tagging, tissue fractionation and enrichment of Aha-labeled ECM proteins were combined with LC-MS/MS analysis. (**B, C**) Relative percentage of matrisome intensity between unenriched (left) and enriched (right) samples. Data points are the average of *n*=3 biological replicates. Labels on the left indicate ECM proteins of interest that were only identified in unenriched samples. Lines connect proteins identified in both unenriched and enriched samples (labeled on the right). Darker, bolded lines highlight ECM proteins of interest and * indicates a significant change (*p*<0.05) in intensity percentage between unenriched and enriched samples. (**D**) Comparison of identified newly synthesized ECM proteins, both unique and shared, between E13.5-E13.75 and E13.5-E14.5 timepoints. (**E**) Comparison of identified newly synthesized ECM proteins, both unique and shared, between P35-P35.25 and P35-P36 timepoints. (**F**) Newly synthesized ECM proteins, both unique and shared, identified in embryonic and adolescent forelimbs.

The coverage of matrisome proteins in the unenriched fractions was comparable to the prior analysis of WT E14.5 and P35 forelimbs (**Tables S3, S5**). Interestingly, Pearson correlation coefficients revealed that E13.75 and E14.5 unenriched samples were less correlated than those from P35.25 and P36, indicating that the matrisome of the adolescent timepoints was less variable than that of the embryos (**Table S5)**. NSPs isolated from the enriched samples were defined as being either (1) exclusively identified in Aha-labeled samples or (2) >2 fold-change in the raw intensity in Aha-labeled compared to PBS samples and statistically significant (*p*≤0.05, two-tailed t-test) (**Figures 5, S7, Table S5**).

We previously showed that the maximum amount of Aha-tagged proteins in the IN fraction occurs around 6-hpi (Saleh, et al., 2019a). Therefore, to identify trends of protein synthesis, the relative percentage of matrisome components was compared between the unenriched and enriched samples 6 hours after Aha administration (**Figure 5B, C)**. Many NSPs followed expected trends based on static matrisome changes during development (**Figure 3A**). For example, there was a significantly higher percentage of COL1A1 in the enriched compared to the unenriched matrisome at E13.75 (**Figure 5B**); however, there was not a difference at P35 (**Figure 5C**). This corresponded with the rapid increase of type I collagen at early timepoints (**Figure 3A**) and indicated that the deposition of fibrillar collagen was approaching homeostasis at P35.25. FN1 and EMILIN1 were found at a significantly lower percentage in the E13.75 enriched sample compared to unenriched and FBN2 was only in the unenriched, indicating that synthesis of FN1, EMILIN1 and FBN2 was decreasing at this timepoint. Of those three proteins, only FN1 was found at P35.25 in the unenriched fraction, supporting our results that all three proteins decreased in abundance over development (**Figure 3A, 5C**). The relative percentage of COL2A1, a fibrillar collagen predominantly found in cartilage (Aszodi, et al., 2001), significantly increased in the E13.75 enriched sample, which correlates with the establishment of the cartilaginous template for the skeletal elements during this time (Martin, 1990). In contrast, the percentage of COL2A1 that was newly synthesized was significantly lower at P35.25, a time when most of the skeleton has undergone endochondral ossification (Karsenty and Wagner, 2002). The relative percentages of COL5A1, COL5A2 and TNC transiently decreased in abundance from E13.5 – E14.5, as well as P3 – P35 (**Figure 3A**), which correlates with the trends observed here.

The distribution of some NSPs was inconsistent with the static proteome. In particular, the relative amount of newly synthesized COL1A2 was not significantly higher than the unenriched at E13.75, unlike what was observed for COL1A1 (**Figure 5B**). This may indicate that the synthesis/degradation rate of the two chains are different. However, it is important to note that the metabolism of Aha *in vivo* is unknown; therefore, it is not clear how long Aha is available for incorporation into NSPs. Alternatively, some NSPs may differentially become cross-linked into the minor part of the matrisome that could not be solubilized prior to enrichment despite the denaturing conditions (Naba, et al., 2017b). Nevertheless, by comparing the NSPs identified at 6- and 24-hpi, biologically-relevant trends were observed. Cartilage-associated proteins (types IX and XI collagen, MATN1) were identified as NSPs exclusively in the embryonic forelimbs (**Figure 5D, 5F**). In addition, the NSPs of adolescent forelimbs corroborated our findings from the static matrisome that CILP2, NID1, TNXB, FMOD, PRELP and LUM, were synthesized and identified only postnatally (P3 and/or P35) (**Figures 3A, 5E, 5F**).

Furthermore, analysis of NSPs between E13.5-E14.5 enabled identification of ECM dynamics that were not revealed by the static proteome. For example, LAMA5 was identified in the enriched proteins of the E13.5-E14.5 labeling window but not identified in the unenriched samples or E14.5 static proteome, indicating this technique could capture and identify NSPs that are in lower abundance (**Figures 3, 5D, 5F Table S5**). The presence of LAMA5 is critical for digit septation; embryos lacking this laminin developed syndactyly, most likely due to defective basement membrane formation that allowed for mesenchymal cells to migrate to the outside of the limb, instead of residing within the interdigit region (Miner, et al., 1998).

Collectively, enrichment of NSPs at the embryonic and adolescent timepoints revealed that different biological processes were occurring during tissue morphogenesis versus growth. Additionally, this technique will enable the identification of ECM proteins that are of low abundance, but nevertheless important for patterning and morphogenesis.

### Comparative analysis of matrisome composition between pathological and WT tissues: osteogenesis imperfecta murine (OIM) forelimbs

To validate that tissue fractionation and LC-MS/MS can resolve differences in matrisome composition as a function of disease or phenotype, CS and IN fractions of E14.5 forelimbs from *osteogenesis imperfecta murine* (OIM, *Col1a2*^OIM^) and WT littermates (*Col1a2*^WT^) were compared (**Figure 6A**). Across all samples, 2211 proteins were identified including 151 ECM proteins (**Table S6**).

**Figure 6.**
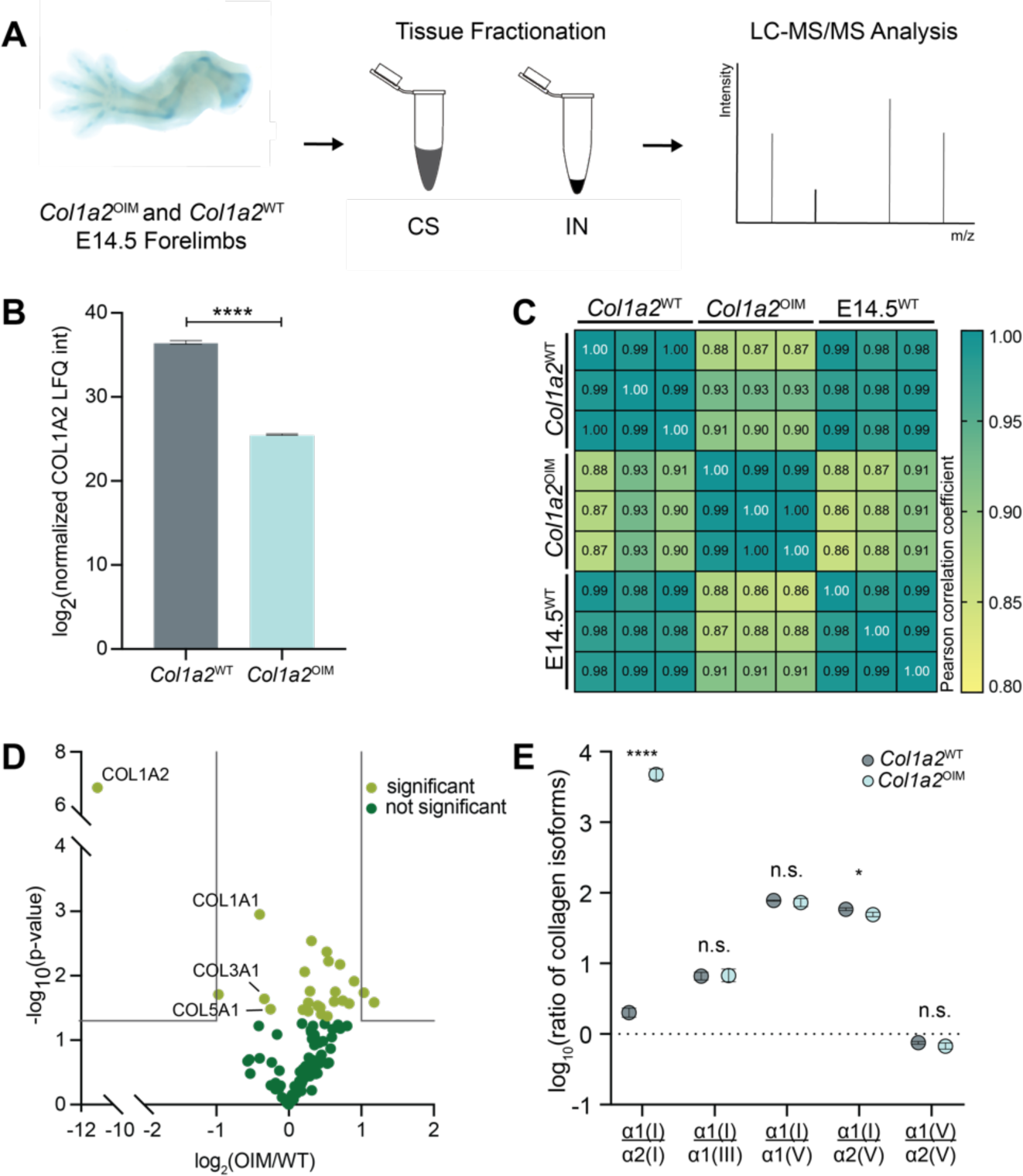
Proteomic analysis of E14.5 *osteogenesis imperfecta murine* (OIM) forelimbs revealed matrisome composition was disrupted. **(A)** Forelimbs of OIM homozygous mutants (*Col1a2*^OIM^) and WT littermates (*Col1a2*^WT^) were fractionated and CS and IN fractions were analyzed by LC-MS/MS (*n*=3 biological replicates). **(B)** Validation of decreased COL1A2 abundance by LC-MS/MS in the IN fractions of *Col1a2*^OIM^ and *Col1a2*^WT^ forelimbs (two-tailed t-test; \*\*\*\**p*<0.0001). **(C)** Pearson correlation coefficients comparing matrisome compositions of the IN fractions of *Col1a2*^OIM^ and *Col1a2*^WT^ forelimbs, as well as E14.5 WT forelimbs (E14.5^WT^) from **Figures 2** and **3**. *Col1a2*^OIM^ and *Col1a2*^WT^ were moderately correlated, whereas *Col1a2*^WT^ and E14.5^WT^ were highly correlated, confirming the reproducibility of the methodology. **(D)** Volcano plot comparing ECM proteins identified in the IN fraction of *Col1a2*^OIM^ and *Col1a2*^WT^ forelimbs. Grey lines denote ≥2-fold change and *p*<0.05 (two-tailed t-test). **(E)** The ratios of collagen chains associated with type I collagen fibrillogenesis in *Col1a2*^OIM^ and *Col1a2*^WT^ forelimbs. Although the ratios between α1(I):α2(I) and α1(I):α2(V) were significantly different in *Col1a2*^OIM^ forelimbs (\**p*<0.05, two-tailed t-test), the other ratios were not affected by the mutation.

In the OIM model, a single nucleotide deletion within the *Col1a2* gene alters the final 50 amino acids of the propeptide (Chipman, et al., 1993). This defect inhibits incorporation of the pro-α2 chains into the type I collagen triple helix, leading to decreased type I collagen fiber content and stability (Weis, et al., 2000; Chipman, et al., 1993). Correspondingly, there was a significant decrease in COL1A2 in OIM mutants (**Figure 6B**). Given that COL1A2 is not incorporated into type I collagen fibrils in the *Col1a2*^OIM^ embryos, we expected this chain would be more soluble compared to COL1A1. Indeed, we found that more COL1A2 was extracted in the CS fraction than COL1A1 (**Table S6**).

Comparison of Pearson correlation coefficients revealed that *Col1a2*^OIM^ and *Col1a2*^WT^ were moderately correlated, suggesting subtle changes in overall matrisome composition as a result of the absence of functional COL1A2 (**Figure 6C**). WT E14.5 forelimbs (E14.5^WT^, **Figure 2**) and *Col1a2*^WT^ showed high correlation, demonstrating the consistency of tissue fractionation across independent experiments for comparing the matrisome of developing tissues.

The abundance of collagens associated with type I collagen fibrillogenesis (types I, III and V), decreased in *Col1a2*^OIM^ forelimbs (**Figure 6D**). Classic phenotypes of osteogenesis imperfecta in humans include increased bone fragility and decreased bone mass due to the significant reduction of type I collagen integration (Morello, 2018). Interestingly, α1(I):α1(III) and α1(I):α1(V) ratios were not affected by the mutation, indicating that absence of COL1A2 does not directly interfere with collagen chain accumulation during fibrillogenesis (**Figure 6E**). During fibrillogenesis, various ECM proteins modify or chaperone collagen precursors and fibrils, including PCOLCE, PLODs and SERPINH1, all of which did not change as a result of the COL1A2 mutation (**Table S6**) (Qi and Xu, 2018; Widmer, et al., 2012; Steiglitz, et al., 2006). Other collagen-modifying proteins, including lysyl oxidase homologs (LOXL1-4), increased in abundance in mutants (**Table S6**). Since the abundance of collagen modifiers in *Col1a2*^OIM^ mutants did not decrease in parallel with type I, III and V collagens, the collagen precursors may be over-modified. Premature crosslinking can halt fibril polymer assembly, leading to the reduced fiber diameter and decreased mechanical strength observed in *Col1a2*^OIM^ mutants (Weis, et al., 2000; Bailey, et al., 1998). Additional studies are needed to validate this hypothesis and explain the decrease in types I, III and V collagens. Nevertheless, these techniques were able to reveal changes in composition and abundance of ECM proteins in a mouse model in which there is a defect in matrix synthesis.

### Comparative analysis of tissue-specific matrisome composition: brains and forelimbs

To further corroborate that these techniques are suitable for comparative developmental proteomics, the matrisome of embryonic brain was compared to that of the forelimb. Brains were collected from WT E14.5 embryos and processed the same as the forelimbs (**Figure 7A**). Across both brain and forelimb tissues, 1873 proteins were identified, 109 of which belonged to the matrisome (**Table S7**). Similar to other embryonic tissues, significantly more matrisome content was identified in the IN fraction (**Figure 7B**). Further, 27 and 33 ECM proteins were identified exclusively in brain and forelimb samples, respectively, and 49 matrisome proteins were identified in both tissues (**Table S7**).

**Figure 7.**
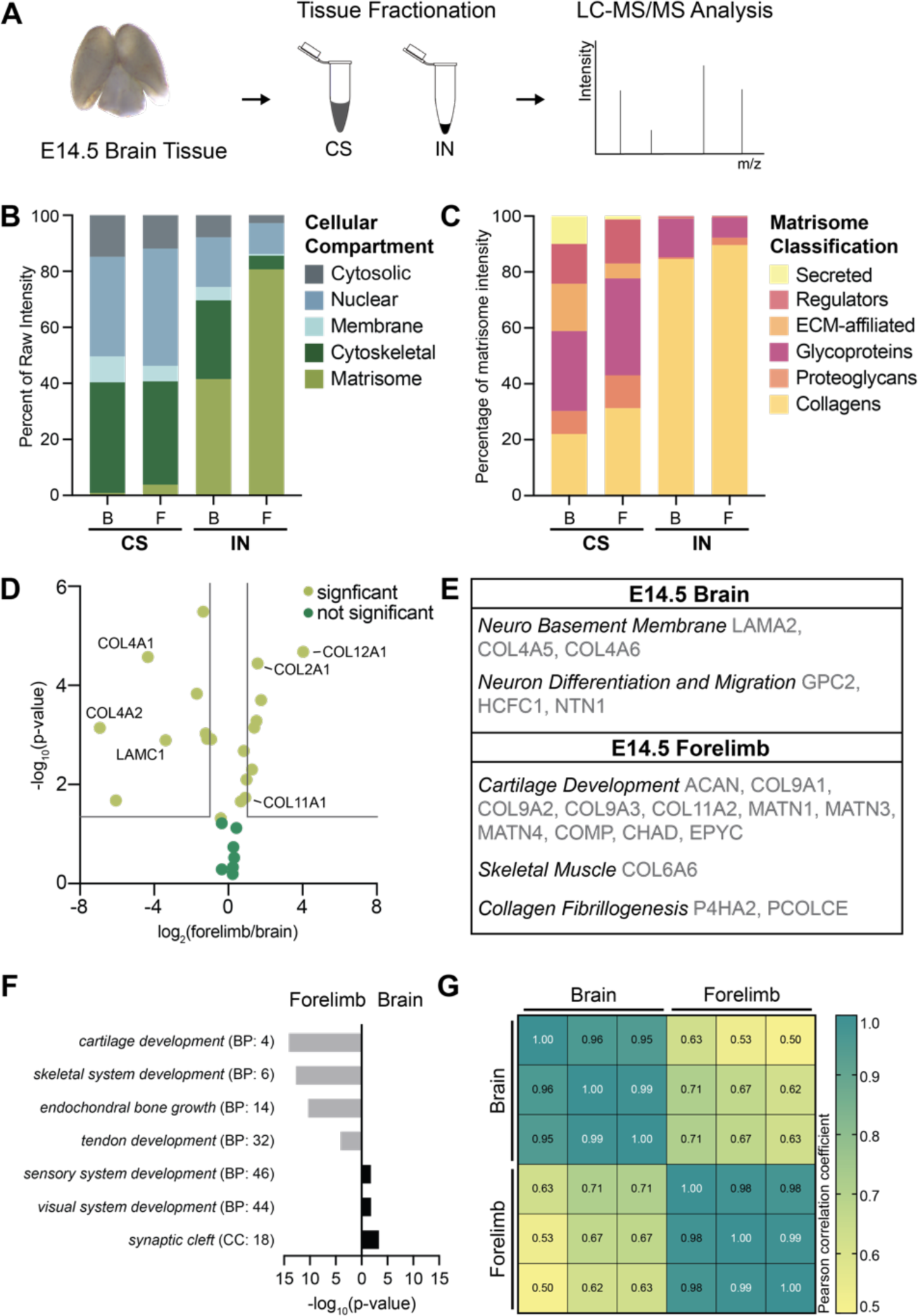
LC-MS/MS analysis demonstrates the matrisome of the embryonic murine brains and forelimbs were significantly different. **(A)** LC-MS/MS analysis of CS and IN fractions of E14.5 brain tissue (*n*=3 biological replicates). **(B)** Proteins annotated by cellular compartment and plotted as average percentage of total raw intensity across biological replicates. B = brain; F = forelimb. **(C)** The distribution of matrisome classifications, determined by percentage of matrisome raw intensity, was significantly different between tissues (*p*<0.0001, two-way ANOVA). **(D)** Volcano plot of ECM proteins identified in the IN fraction of brain and forelimb tissues. Grey lines denote ≥2-fold change and *p*<0.05. **(E)** List of select ECM proteins, exclusively identified in the brain or forelimb, grouped by biological function to highlight distinct matrisome components of each tissue. **(F)** Select GO terms associated with ECM proteins more abundant or exclusively identified in forelimb and brain tissues (**Table S7**). **(G)** Comparison of Pearson correlation coefficients revealed differences in the matrisome identified in the IN fractions of brain and forelimb tissues.

Proteins associated with cartilage (COL2A1, COL11A1) and tendon (COL12A1) (Zou, et al., 2014) were significantly more abundant in forelimbs, whereas some basement membrane proteins were higher in brain (COL4A1, COL4A2, LAMC1) (**Figure 7D**). COL4A5 and COL4A6 were identified exclusively in brain, which are present as α5(IV)2α6(IV) heterotrimers in the pia mater basement membrane (Hubert, et al., 2009). Other ECM proteins critical for formation and maturation of the blood brain barrier (AGRN) (Barber and Lieth, 1997) were exclusively identified in the brain (**Figure 7E, Table S7**); whereas, developing cartilage (type IX collagen, matrilins, COMP, CHAD) and skeletal muscle (COL6A6) matrisome constituents were only found in the forelimb (Ocken, et al., 2020; Acharya, et al., 2014; Hessle, et al., 2013; Sabatelli, et al., 2012). GO analysis of ECM proteins that were more abundant in, or exclusive to, brain or forelimb tissues generated neuro- or musculoskeletal-related GO terms, respectively (**Figure 7F**). Further, there was low correlation between the forelimb and brain matrisome (**Figure 7G**), suggesting early differentiation of the ECM by tissue type and function. Taken all together, these results indicated tissue fractionation combined with LC-MS/MS was able to resolve tissue-specific differences in ECM composition during development.

Until now, an open question in the field of limb and musculoskeletal development was the contribution of the ECM during tissue assembly. Results from our study provide a resource that describes how the matrisome changes during forelimb development. In addition, we show how modifications of existing methods enable the identification of the static and newly synthesized matrisome in the developing forelimb. As the ECM is a critical component of all organ systems, this information will be a valuable guide for future investigations into the roles of ECM proteins during morphogenesis and growth.

## Acknowledgements

The authors would like to thank Dr. Uma Aryal and Victoria Hedrick of the Purdue Proteomics Core and members of the Calve and Kinzer-Ursem laboratories for helpful comments and discussion. This work was supported by the National Institutes of Health [R21 AR069248 and R01 AR071359 to S.C. and T.L.K.; DP2 AT009833 to S.C.].

## Author Contributions

Conceptualization, K.R.J and S.C.; Methodology, K.R.J., A.M.S., S.N.L., A.R.O., T.L.K, and S.C.; Validation, K.R.J, S.N.L., and S.C.; Investigation, K.R.J., A.M.S., and S.N.L.; Formal Analysis, K.R.J., A.M.S., A.R.O., and S.C.; Writing – Original Draft, K.R.J., S.N.L., and S.C.; Writing – Review & Editing, K.R.J., A.M.S., S.N.L., A.R.O., T.L.K, and S.C.; Visualization, K.R.J, S.N.L, and S.C.; Supervision, T.L.K. and S.C.; Project Administration, K.R.J., T.L.K., and S.C.; Funding Acquisition, T.L.K. and S.C.

## STAR Methods

### KEY RESOURCES TABLE

| REAGENT or RESOURCE | SOURCE | IDENTIFIER |
| --- | --- | --- |
| **Antibodies** (dilutions can be found in **Table S4**) |  |  |
| Rabbit polyclonal anti-type I collagen | Millipore | Cat# AB765P<br>RRID: AB_92259 |
| Goat polyclonal anti-type V collagen | Southern Biotech | Cat# 1350-01<br>RRID: AB_2794740 |
| Mouse IgG1 monoclonal anti-skeletal muscle myosin heavy chain (MY32; clone MYSN02) | Thermo Fisher Scientific | Cat# MA5-11748<br>RRID: AB_10979315 |
| Rat monoclonal anti-tenascin C (TNC; clone 578) | R & D Systems | Cat# MAB2138<br>RRID: AB_2203818 |
| Rabbit polyclonal anti-fibrillin 2 (FBN2) | Gift from Dr. Robert Mecham | (Beene, et al., 2013) |
| Mouse IgG1κ monoclonal anti-nidogen-2 (NID2, clone F-2) | Santa Cruz Biotechnology | Cat# sc-377424<br>RRID: AB_2819357 |
| Rabbit polyclonal anti-EMILIN1 | Novus Biologicals | Cat# NBP1-84127<br>RRID: AB_11037041 |
| Rat monoclonal perlecan (HSPG2; clone A7L6) | Santa Cruz Biotechnology | Cat# sc-33707<br>RRID: AB_627714 |
| Alexa Fluor 488-conjugated goat anti-rabbit secondary antibody | Thermo Fisher Scientific | Cat# A-11034<br>RRID: AB_2576217 |
| Alexa Fluor 488-conjugated donkey anti-rat secondary antibody | Thermo Fisher Scientific | Cat# A-21208<br>RRID: AB_2535794 |
| Alexa Fluor 546-conjugated goat anti-mouse IgG1 secondary antibody | Thermo Fisher Scientific | Cat# A-21123<br>RRID: AB_2535765 |
| Alexa Fluor 555-conjugated donkey anti-goat secondary antibody | Thermo Fisher Scientific | Cat# A-21432<br>RRID: AB_2535853 |
| Alexa Fluor 633-conjugated goat anti-mouse IgG1 secondary antibody | Thermo Fisher Scientific | Cat# A-21126<br>RRID: AB_2535768 |
| Alexa Fluor 647-conjugated donkey anti-rabbit secondary antibody | Thermo Fisher Scientific | Cat# A-31573<br>RRID: AB_2536183 |
| **Deposited Data** |  |  |
| Mass spectrometry proteomics data | This paper | MassIVE: MSV000085556 |
| **Experimental Models: Organisms/Strains** |  |  |
| *Mus musculus*: C57BL/6J (WT) | The Jackson Laboratory | Stock #: 000664 |
| *Mus musculus*: B6C3Fe a/a-*Col1a2<sup>oIM</sup>/J*<br>(COL1A2) | The Jackson Laboratory | Stock #: 001815 |
| **Software** |  |  |
| Fiji Software | (Schindelin, et al., 2012) |  |
| Illustrator | Adobe | v. 2019 |
| Photoshop | Adobe | v. 2019 |
| Excel | Microsoft | v. 2020 |
| Prism | GraphPad | v. 8.3.1 |
| Perseus | (Tyanova, et al., 2016) |  |
| MaxQuant | (Cox and Mann, 2008) | v. 1.6.1.0 |
| **Chemicals, Peptides and Recombinant Proteins** |  |  |
| L-azidohomoalanine | Click Chemistry Tools | Catalog #: 1066 |
| Halt Protease Inhibitor Cocktail (100×) | Thermo Fisher Scientific | Catalog #: 78429 |
| Chondroitinase ABC from *Proteus vulgaris* | Sigma-Aldrich | Catalog #: C3667 |
| Diazo Biotin Alkyne | Click Chemistry Tools | Catalog #: 1042 |
| THPTA (tris(3-hydroxypropyltriazolylmethyl)amine) | Click Chemistry Tools | Catalog #: 1010 |
| Pierce NeutrAvidin Agarose | Thermo Fisher Scientific | Catalog #: 29200 |
| Endoproteinase LysC | New England Biolabs | Catalog #: P8109S |
| Pierce Trypsin Protease, MS Grade | Thermo Fisher Scientific | Catalog #: 90057 |
| Pierce Detergent Removal Spin Columns | Thermo Fisher Scientific | Catalog #: 87777 |
| C-18 MicroSpin columns | The Nest Group, Inc. | Catalog #: SS18V |
| Mouse on Mouse (MOM) Basic Kit | Vector Laboratories | Catalog #: BMK-2202 |
| ImmEdge pen | Vector Laboratories | Catalog #: H-4000 |
| Fluoromount G | SouthernBiotech | Catalog #: 0100-01 |
| **Critical Commercial Assays** |  |  |
| Compartment Protein Extraction Kit | Millipore EMD | Catalog #: 2145 |
| Pierce 660 nm Protein Assay | Thermo Fisher Scientific | Catalog #: 22662 |
| Pierce Quantitative Colorimetric Peptide Assay | Thermo Fisher Scientific | Catalog #: 23275 |
| **Other** |  |  |
| TissueRuptor | Qiagen | https://www.qiagen.com/us |
| CentriVap Benchtop Vacuum Concentrator | Labconco | https://www.labconco.com/ |
| Branson Ultrasonics Sonifier S-450A | Fisher Scientific | Catalog #: 22-309783 |
| ThermoMixer, F1.5 | Eppendorf | Catalog #: 5384000020 |
| Q Exactive™ HF Hybrid Quadrupole-Orbitrap Mass Spectrometer | Thermo Fisher Scientific | Catalog #: IQLAAEGAAPFAALGMBFZ |
| Shandon Cryotome FE | Thermo Fisher Scientific | Catalog #: A78910100 |
| Leica DMI6000 Microscope | Leica Microsystems | https://www.leica-microsystems.com/ |

### RESOURCE AVAILABILITY

#### Lead Contact

Further information and requests for resources and reagents should be directed to and will be fulfilled by the Lead Contact, Sarah Calve.

#### Materials Availability

This study did not generate new unique reagents or mouse lines.

#### Data and Code Availability

The RAW data files generated during this study are available at MassIVE, MSV000085556.

### EXPERIMENTAL MODEL AND SUBJECT DETAILS

All experimental protocols were performed in accordance with the guidelines established by the Purdue Animal Care and Use Committee, and all methods were approved by this committee (PACUC; protocol# 1209000723). PACUC ensures that all animal programs, procedures, and facilities at Purdue University adhere to the policies, recommendations, guidelines, and regulations of the USDA and the United States Public Health Service in accordance with the Animal Welfare Act and Purdue’s Animal Welfare Assurance.

Wild-type (WT) C57BL/6J and B6C3Fe *a/a-Col1a2*^*OIM*^*/J* (COL1A2)(Chipman, et al., 1993) mice were obtained from The Jackson Laboratory. For all *in vivo* studies (timed-mating), the sex of the embryos was not determined prior to analysis. Female mice were time-mated, where noon of the day a copulation plug was found was considered to be embryonic day (E)0.5. All mice were euthanized via CO_2_ inhalation, followed by cervical dislocation, with the exception of post-natal day (P)3 pups in which decapitation was used.

### METHOD DETAILS

#### Experimental Groups: Tissue Collection

#### WT whole embryos

Embryos were collected at embryonic day (E) 11.5 through E14.5, weighed (see **Tables S1, S2**), snap frozen, and stored at -80°C.

#### WT forelimbs

Mouse forelimbs were microdissected away from embryos, taking care to maintain the musculoskeletal structures from the scapula to the digit tips. Forelimbs were isolated from E11.5-E14.5 embryos, P3 pups and P35 mice, weighed (see **Table S3**), snap frozen and stored at -80°C. Forelimbs were predominantly comprised of developing musculoskeletal tissue with the exception of the skin. For E11.5-E14.5, the skin was left on the forelimbs since it made up less than 1% of the wet weight and was challenging to remove while keeping the musculoskeletal tissues intact. The skin was removed from P3 and P35 forelimbs prior to analysis since it was ∼25% of the total wet weight of the tissue and easier to remove (data not shown). Skin and hair were removed from P3 and P35 forelimbs.

#### Aha-labeled WT forelimbs

The methionine analog L-azidohomoalanine (Aha) was reconstituted in PBS and adjusted to pH 7.4 with NaOH to generate a final stock concentration of 10 mg/mL. Aha stock solutions were sterilized using 0.2 μm cellulose acetate membrane filters and stored at -20°C.

For embryonic forelimb tissues, WT pregnant dams were injected subcutaneously at the base of the neck, to avoid piercing the amniotic sac, with Aha (0.1 mg Aha/g mouse) at E13.5 and sterile PBS pH 7.4 was used for control injections (10 µL PBS/g mouse). Dams were euthanized and forelimbs were collected as described above at either E13.75 (6 hours post-injection; hpi) or E14.5 (24 hpi). To collect adolescent forelimbs, non-pregnant WT female mice were injected at P35 and forelimbs were harvested at P35.25 (6 hours post-injection) or P36 (24 hours post-injection) as described above. Tissues were weighed (see **Table S5**), snap frozen and stored at -80°C.

#### COL1A2 OIM forelimbs

Heterozygous COL1A2 (COL1A2^+/-^) male and female mice were time-mated to obtain COL1A2^+/+^ (*Col1a2*^WT^) and COL1A2^-/-^ (*Col1a2*^OIM^) embryos. Forelimbs were microdissected away from the embryos at E14.5, weighed (see **Table S6**), snap frozen and stored at -80°C.

#### WT brains

Mouse brains, including both forebrain and hindbrain, were microdissected away from E14.5 embryos, weighed (see **Table S7**), snap frozen and stored at -80°C.

Details regarding the biological replicates of each experimental group for tissue fractionation and proteomic analysis are listed in **Tables S1-3, 5-7**.

#### Tissue Fractionation

Proteins were extracted from tissues using buffers from the Compartment Protein Extraction Kit as previously described (Naba, et al., 2015). Briefly, tissues were homogenized (TissueRuptor) in ice-cold C (cytosolic) buffer and rotated end-over-end for 30 minutes at 4°C. Buffer C, as well as all subsequent buffers also contained protease inhibitors. Aliquots of E12.5 whole embryo homogenate were collected after initial homogenization, snap frozen and stored at -80°C. Following incubation, samples were centrifuged for 20 minutes at 16,000 × *g*. Supernatants were collected (C fraction), snap frozen and stored at -80°C. Pellets were resuspended with W (wash) buffer and washed by end-over-end rotation at 4°C for 5 minutes. Samples were centrifuged and supernatants were collected, snap frozen and stored at -80°C. Subsequent incubations with the N (nuclear) buffer supplemented with 0.1% benzonase (2×) and the M (membrane) buffer (1×), were 30 minutes at 4°C prior to centrifugation and fraction collection (N and M fractions, respectively). The final incubation with CS (cytoskeletal) buffer was performed at room temperature for 20 minutes prior to centrifugation and CS fraction collection. The remaining pellet was considered the insoluble fraction (IN), and was washed 3× with PBS, snap frozen, and stored at -80°C.

#### Protein Quantification of Fractionated Forelimbs

To quantify the amount of protein extracted by each tissue fractionation buffer, another set of E11.5 – E14.5, P3 and P35 forelimbs were collected and fractionated as described above. IN fractions were resuspended in 8M urea/100mM ammonium bicarbonate. Protein concentration for each fraction was measured using the Pierce 660 nm Quantitative Colorimetric Assay and amount of protein in each fraction per forelimb was calculated using **Eq. 1**

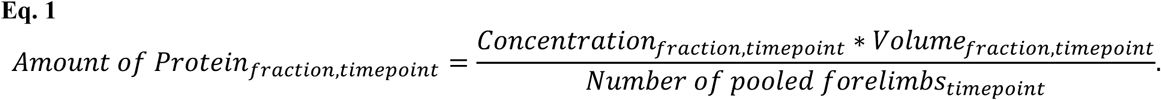

The total amount of protein in each forelimb (**Figure S3B**) was determined by summation of the amount of protein in all fractions and normalized to the number of forelimbs in each biological replicate.

#### Enrichment of Aha-labeled ECM Proteins

To identify the newly synthesized proteins at embryonic and adolescent timepoints, forelimbs labeled with Aha were fractionated as described above with slight modifications in buffer volumes (**Table S5**). IN fractions were resuspended in 700μL of Chondroitinase ABC Digestion Buffer (0.1M Tris-HCl, 0.03M sodium acetate, pH 8) with Halt protease inhibitor cocktail (final concentration 1×). Chondroitinase ABC was added to each sample (final concentration 0.2U/700μL) and incubated overnight at 37°C with constant agitation (1000 rpm, ThermoMixer). After incubation, four volumes of 100% acetone were added to each sample and incubated at -20°C to precipitate proteins. Proteins were pelleted by centrifugation for 20 minutes at 4°C, acetone was removed, and pellets were vacuum dried (Centrivap). Dried pellets were resuspended in 500μL 8M urea/100mM ammonium bicarbonate and sonicated on ice 4 × 10 seconds using 50% duty cycle and output 3 (Sonifier). Samples were centrifuged at 16,000 × *g* for 20 minutes, the supernatant was transferred to a new tube and protein concentration was measured using the Pierce 660 nm Quantitative Colorimetric Assay. Aliquots of each supernatant were transferred to a new tube, labeled as the “Unenriched” fraction for each sample, snap frozen and stored at -80°C until protein digestion.

Aha-labeled proteins were enriched as described by (Saleh, et al., 2019a) with slight modifications. Proteins were first alkylated with iodoacetamide (IAA, final concentration, 39mM) for 30 minutes at RT, protected from light. After alkylation, click reagents were added individually in the following order: (a) cleavable diazo biotin-alkyne probe (DBA, final concentration 0.05mM); (b) ligand tris(3-hydroxypropyltriazolylmethyl)amine (final concentration 10mM); (c) copper sulfate (final concentration 2mM); (d) aminoguanidine (final concentration 20mM); and (e) sodium ascorbate (final concentration 5mM). Click reactions were rotated end-over-end for 3 hours at room temperature, followed by protein precipitation with acetone.

Protein pellets were resuspended in 550μL 4.4M urea/100mM ammonium bicarbonate buffer and remaining precipitates were pelleted by centrifugation at 16,000 × *g* for 20 minutes at room temperature. Supernatants were added to 200μL of 50% NeutrAvidin bead slurry, previously washed 3 × 1mL of 100mM ammonium bicarbonate, and rotated end-over-end for 1.5 hours at room temperature. After incubation, beads were washed 4 × 10 minutes with 1mL of 4M urea and 0.1% sodium dodecyl sulfate (SDS) in 100mM ammonium bicarbonate and 3× for 10 min with 1mL of with 0.1% SDS in PBS (pH 7.4).

Bound proteins were eluted by resuspending the beads in 400μL 50mM sodium dithionite with 0.1% SDS in PBS (pH 7.2) and rotating end-over-end for 1 hour at room temperature protected from light. The elution fraction was collected by centrifugation for 2 minutes at 1,200 × *g.* The elution step was repeated two more times and fractions were combined for the “Enriched” sample. Enriched proteins were precipitated with acetone, snap frozen and stored at -80°C.

Enriched and Unenriched samples were processed for LC-MS/MS analysis as delineated above. The peptide concentrations of the Enriched samples were normalized such that the most concentrated Aha sample of each condition (E13.75, E14.5, P35.25, P36) was diluted to 0.2 mg/mL and equivalent volumes were added to the remaining Aha and PBS samples. All unenriched samples were normalized to 0.2 mg/mL.

#### Liquid Chromatography Tandem Mass Spectrometry (LC-MS/MS) Analysis

##### Fractions analyzed for each experimental group

For E12.5 whole embryos, homogenates, C, N, M, CS and IN fractions were analyzed by LC-MS/MS. For E11.5 – E14.5 whole embryos C, N and M fractions were combined into one CNM sample before LC-MS/MS analysis. CNM, CS and IN fractions were analyzed by LC-MS/MS. For E11.5 – E14.5, P3 and P35 forelimbs, CS and IN fractions were analyzed by LC-MS/MS. For Aha-labeling experiments, enriched and unenriched samples from E13.75, E14.5, P35.25 and P36 timepoints were analyzed by LC-MS/MS. For E14.5 *Col1a2*^WT^ and *Col1a2*^OIM^ forelimbs, CS and IN fractions were analyzed by LC-MS/MS. For E14.5 brains, CS and IN fractions were analyzed by LC-MS/MS.

##### Enzymatic digestion of proteins

IN fractions and enriched samples were resuspended in 8M urea/100mM ammonium bicarbonate and reduced with dithiothreitol (DTT, final concentration 10mM) for 2 hours at 37°C with constant agitation. All other fractions and samples were diluted 1:2 with 8M urea in 100mM ammonium bicarbonate and reduced with DTT. Samples were brought to room temperature prior to alkylation with IAA (final concentration 25mM) for 30 minutes in the dark. Samples were diluted to 2M urea with 100mM ammonium bicarbonate and deglycosylated with 0.1U/200μL chondroitinase ABC for 2 hours at 37°C with constant agitation. Proteins were then digested into peptides by three enzymatic steps at 37°C with constant agitation: (1) endoproteinase LysC (1μg/200μL) for 2 hours; (2) MS-grade trypsin (3μg/200μL) overnight; and (3) MS-grade trypsin (1.5μg/200μL) for an additional 2 hours. Digestion enzymes were inactivated by acidification (trifluoroacetic acid, TFA, final concentration 0.1%).

Detergent contamination was removed from samples derived from M, CS and IN fractions using Pierce Detergent Removal Spin Columns per the manufacturer’s protocol. All samples were desalted using C-18 MicroSpin columns. Briefly, columns were prepared with 100% acetonitrile (ACN) and HPLC-grade water/0.1% TFA. Peptides were then added to the columns and washed with two 100μL volumes of water/0.1% TFA before elution with 50μL of 80% ACN/25mM formic acid (FA). After elution, samples were dried at 45°C for 4 hours and peptides were resuspended in 10μL of 3% ACN/0.1% FA. Peptide concentration was measured using the Pierce 660 nm Quantitative Colorimetric Assay. Peptide concentrations for all fractions were normalized to 1 μg/μL with 3% ACN/0.1% FA.

##### LC-MS/MS

Samples were analyzed using the Dionex UltiMate 3000 RSLC Nano System coupled to the Q Exactive™ HF Hybrid Quadrupole-Orbitrap Mass Spectrometer. Following digestion and clean up, 1μg of peptide was loaded onto a 300μm i.d. × 5mm C18 PepMap™ 100 trap column and washed for 5 minutes using 98% purified water/2% ACN/0.01% FA at a flow rate of 5 μL/minute. After washing, the trap column was switched in-line with a 75 μm × 50 cm reverse phase Acclaim™ C18 PepMap™ 100 analytical column heated to 50°C. Peptides were separated using a 120 minute gradient elution method at a flow rate of 300 nL/minute. Mobile phase A consisted of 0.01% FA in water while mobile phase B consisted of 0.01% FA in 80% ACN. The linear gradient started at 2% B and reached 10% B in 5 minutes, 30% B in 80 minutes, 45% B in 91 minutes, and 100% B in 93 minutes. The column was held at 100% B for the next 5 minutes before being brought back to 2% B and held for 20 minutes. Samples were injected into the QE HF through the Nanospray Flex™ Ion Source fitted with an emission tip (New Objective). Data acquisition was performed monitoring the top 20 precursors at 120,000 resolution with an injection time of 100 ms.

#### Forelimb Tissue Preparation for Immunohistochemistry (IHC)

Microdissected forelimbs from E11.5 – E14.5, P3 and adult (between 6 – 20 weeks) mice were either directly embedded in optimal cutting temperature (OCT) compound or processed as follows based on antibody compatibility (**see Table S4**). Forelimbs were incubated in 4% paraformaldehyde (PFA) in PBS for 2 – 4 hours at room temperature, washed in PBS, and incubated overnight at 4°C in sucrose solution (23 wt/wt % solution of sucrose in PBS with 0.02% sodium azide). Forelimbs were incubated in a 50% OCT:50% sucrose solution for 30 minutes before transferring to a mold containing OCT. Samples were frozen with dry ice-cooled isopentane and stored at -80^°^C. Sections of 10 μm forelimb tissue were acquired using a Shandon Cryotome FE, adhered to charged slides and stored at -20°C.

#### IHC Analysis

Incubations were conducted at room temperature unless indicated otherwise (**Table S4**) and samples were protected from light when fluorescent secondary staining reagents were used. Tissue sections were equilibrated to room temperature, encircled using an ImmEdge pen, rehydrated in PBS for 10-15 minutes, fixed with 4% PFA for 5 minutes and washed in PBS. Sections were permeabilized with 0.1% Trition-100 in PBS for 5 minutes and rinsed briefly with PBS prior to blocking for 1 hour with IgG blocking buffer from the Mouse on Mouse (MOM) Basic Kit following manufacturer’s instructions. Tissues were washed 3 × 2 minutes with PBS and blocked for 5 minutes with protein diluent from the MOM Basic kit. Primary antibodies, in solution with the MOM protein diluent, were applied to tissues for varying times and concentrations (indicated by **Tables S4**), then washed 3 × 2 minutes with PBS.

Secondary antibodies and DAPI for NID2/FBN2/TNC and EMILIN1/HSPG2/MY32 combinations were applied to tissue in a solution of MOM protein diluent for times and concentrations indicated in **Table S4**. Slides were washed 3 × 2 minutes with PBS. For the COL I/COL V/MY32 combination, sequential staining was performed due to incompatibilities between secondary antibodies (**Table S4**). Tissues were incubated in MOM protein diluent for 5 minutes, followed by a solution of donkey anti-goat secondary in MOM protein diluent and DAPI for times and concentrations indicated in **Table S4**. Slides were washed 3 2 minutes with PBS and incubated for 5 minutes with MOM protein diluent. Secondary antibody solution of goat anti-mouse, rabbit, and DAPI were applied to tissue as described in **Table S4**. Slides were washed 3 × 2 minutes with PBS.

#### Imaging Analysis

Coverslips were mounted using FluoromountG and sealed with clear nail polish. Slides were stored at 4°C until imaged with a Leica DMI6000 at 20× magnification. Images were processed and compiled using Fiji Software and Adobe Photoshop, respectively, and exposure settings were consistent between the sample and negative controls for individual timepoints.

### QUANTIFICATION AND STATISTICAL ANALYSIS

When appropriate, statistical tests and number of biological replicates used for each graphical analysis are reported in the figure legends or supplemental tables. Statistical significance was determined by *p*<0.05 for two tailed t-tests or one- or two-way ANOVAs. Error bars and ± values report the standard deviation of the mean. Data processing was conducted with Microsoft Excel (for filtering and data handling), Perseus (for clustering analysis) (Tyanova, et al., 2016), GraphPad Prism 8 (for data visualization and statistical analysis), and Adobe Illustrator/Photoshop (for figure compilation).

#### LC-MS/MS Data Processing

Raw files were analyzed with MaxQuant (Cox and Mann, 2008). Default settings were used unless noted otherwise (**Tables S1-3, 5-7**). Peak lists were searched against the *Mus musculus* UniProt FASTA database (November 2018). In Aha-enrichment of NSPs experiments, peak lists were also searched against the *Gallus gallus* Avidin FASTA protein sequence (May 2018). Match-between-runs was enabled between biological replicates. Cysteine carbamidomethylation was included as a fixed modification and variable modifications included oxidation of methionine, hydroxylysine, hydroxyproline, deamidation of asparagine, and conversion of glutamine to pyro-glutamic acid. In Aha-enrichment of NSPs experiments, two additional modifications were included: Aha substitution for methionine and cleaved DBA-tagged Aha substitution for methionine. Peptide and protein false discovery rates were set to 0.01 and determined by a reverse decoy database derived from the *Mus musculus* database.

#### Proteomic Data Analysis

Proteins that were identified by one unique or razor peptide across all samples, or labeled as a potential contaminant or reverse hit, were filtered from the data set. Further, proteins were removed if an intensity value was only found in one biological replicate within an experimental group. After filtration, proteins were classified into Cellular Compartments (CC): cytosolic, nuclear, membrane, cytoskeletal and matrisome (Saleh, et al., 2019a; Naba, et al., 2012). ECM proteins were further categorized into Matrisome Classifications (MCs): secreted factors, ECM regulators, ECM-affiliated proteins, ECM glycoproteins, proteoglycans and collagens (Naba, et al., 2012).

For whole WT embryos (**Tables S1, S2**), WT forelimbs (**Table S3**), OIM forelimbs (**Table S6**) and WT brain (**Table S7**) analyses, label free quantification (LFQ) was employed to compare protein intensities (LFQ intensities) across samples, while raw intensities were used for intrasample comparisons. In the Aha-enrichment analysis **(Table S5**), raw intensities were used to identify newly synthesized ECM proteins and corresponding data analysis. Raw and LFQ intensities were normalized for each analysis as delineated below. An overview of each proteomic data workflow is shown in **Figure S7**.

#### Cellular compartment and matrisome classification percentages

The percentage of each CC in a sample was calculated by **Eq. 2**,

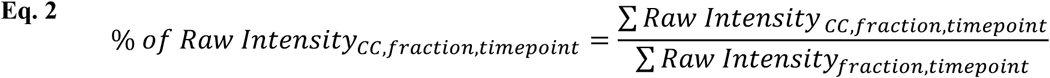

and values were averaged across biological replicates prior to graphical analysis (**Figures S1B, 1B, 2C, 7B**). The distribution of MCs at each timepoint were calculated by **Eq. 3**,

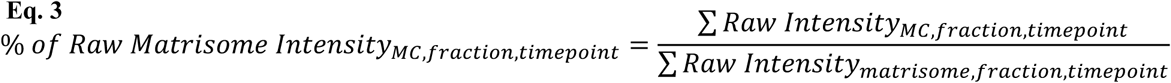

and values were averaged across biological replicates prior to graphical analysis (**Figures 2D, 7C**).

#### Intrasample collagen isoform ratios

Raw intensities (normalization was experiment-specific, see sections below or **Figure S7**) were used for intrasample comparisons (**Figures 3B, 6E**). One-way ANOVA was used to determine significant differences between ratios (**Tables S3, S6**).

#### Volcano plot analysis

For subsequent volcano plot analyses (**Figures 3C, 3D, 6D** and **7D**), LFQ protein intensities (normalization was experiment-specific, see sections below or **Figure S7**) were log_2_ transformed and averaged across biological replicates. The fold change and *p*-values from corresponding two-tailed t-tests were plotted to compare matrisome composition.

#### Whole embryo quantitative analysis

For Gene Ontology (GO) analysis of E14.5 whole embryos (**Figure 1C**), terms associated with the 50 most abundant proteins in the CNM, CS and IN fractions of E14.5 whole embryos were determined using g:Profiler (Raudvere, et al., 2019). The top five enriched “Cellular Component” and “Biological Process” GO terms from each list, along with the -log_10_(*p-value*) were reported. Terms that had a lower ranking in the GO analysis of a specific fraction but were top five in another fraction were also included.

To view enrichment of ECM proteins in E14.5 embryos (**Figure S2**), the same proteins were used in unbiased clustering heat map analysis. Row z-scores for LFQ intensities were calculated across all timepoints and proteins were subjected to unbiased clustering (Euclidean) for heatmap analysis using Perseus.

#### Forelimb quantitative analysis

Many of the ECM proteins identified in E11.5 - E14.5, P3 and P35 forelimbs were present in both CS and IN fractions (**Figure S3A**). To evaluate the effects of combining intensities from CS and IN fractions on observed proteomic trends, LFQ intensities of ECM proteins in the CS and IN fractions were normalized to adjust intensities to reflect equivalent amounts of ECM in each fraction using **Eq. 4**,

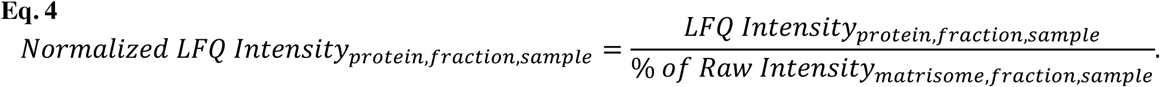

Normalized LFQ intensities for CS and IN samples (**Figure S3C**, top, middle) were used to compare protein trends from combined IN and CS LFQ intensities (**Figure S3C**, bottom).

Matrisome intensities were combined such that the ratio between protein content was maintained. LFQ intensity values were assumed representative of 1 μg of sample was loaded; therefore, to combine individual protein intensities from CS and IN fractions, intensities were scaled to reflect the total amount of protein in that fraction by **Eq. 5**.

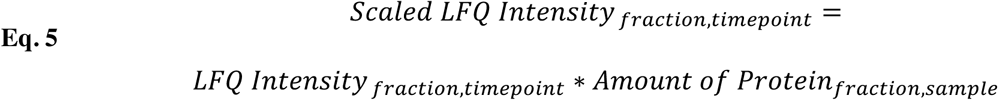

LFQ intensities were then added using **Eq. 6**, with assumption that the scaled LFQ signal is additive based on the peptides compared,

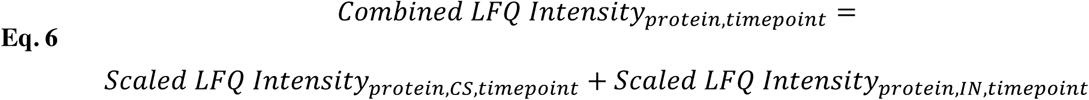

Combined LFQ protein intensities were then normalized to be representative of equivalent amounts of ECM across timepoints by **Eq. 7**,

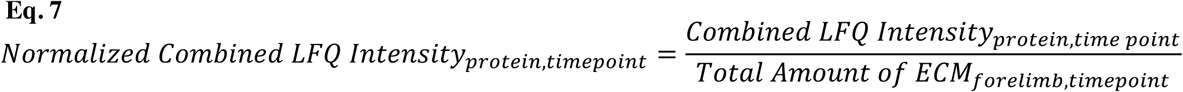

The total amount of ECM per forelimb (**Figure S3B**) was determined using the amount of protein (calculated by **Eq. 1**) and the percentage of matrisome in the CS and IN fractions (calculated by **Eq. 2**), shown in **Eq. 8**,

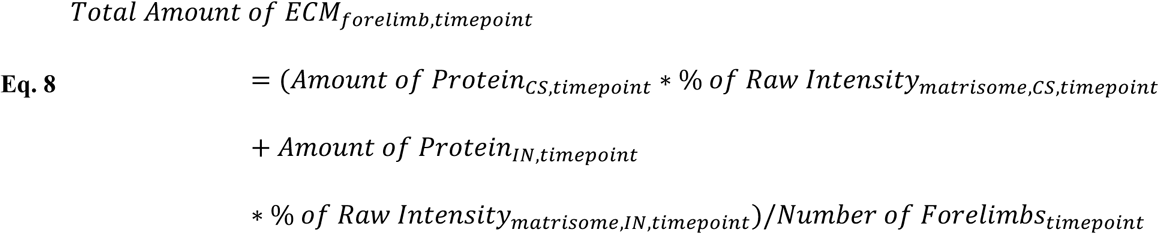

To resolve the relative amounts of ECM protein intensity in CS and IN fractions, separately, additional heat maps were generated using -log_10_ transformed scaled LFQ intensities (**Eq. 5, Figure S3E**, top and middle). The percentage of combined LFQ intensity (**Figure S3E**, bottom) attributed to the IN fraction was plotted as a heat map (**Figure S3D**) to reveal which matrisome classifications were more prominent in the IN fraction.

Normalized combined LFQ intensities were used for subsequent WT forelimb graphical and statistical analyses, unless otherwise noted (see **Figure S7**).

To visualize matrisome dynamics as a function of development, row z-scores were calculated for each ECM protein and averaged across biological replicates. ECM proteins were arranged in a heat map (**Figure 3A**) to show protein dynamics during morphogenesis and growth. Volcano plot analysis (**Figures 3B, C**) and Pearson correlation coefficients heat maps (**Figure 3D**) were used to further ascertain differences in ECM composition between timepoints.

To determine the abundance ratios between specific collagen isoforms (**Figure 3E**), raw intensities were combined and normalized, as delineated above for LFQ intensities (**Eqs. 2, 4-8, Figure S7**), and used. Ratios were log_10_ transformed, averaged across biological replicates and plotted.

#### Aha-enriched ECM proteins quantitative analysis

The newly synthesized matrisome was identified as ECM proteins that were (1) exclusive to Aha-labeled samples, or (2) the fold change of raw intensity in Aha-labeled samples, compared to negative (PBS) control, was >2 and *p* < 0.05. The relative percentage of matrisome intensity for each ECM protein was calculated for unenriched and enriched samples by **Eq. 9**,

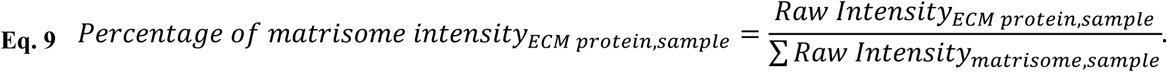

Relative percentages in enriched samples were compared to unenriched using two-sided t-tests where *p* < 0.05 was significant. Percentages for unenriched and enriched samples were plotted as an average across biological replicates (**Figure 5B, C).**

#### COL1A2 OIM forelimbs quantitative analysis

Previous analysis of WT forelimbs revealed minimal changes in protein trends after combining CS and IN intensities; therefore, we analyzed CS and IN fractions separately for comparative analysis of *Col1a2*^WT^ and *Col1a2*^OIM^ forelimbs. After protein filtration, LFQ intensities of ECM proteins in the CS and IN fractions were normalized using **Eq. 2** and **Eq. 4**. Normalized LFQ intensities were used for subsequent analysis, unless otherwise noted. Pearson correlation coefficient analysis was conducted to determine the degree of similarity between matrisome composition of *Col1a2*^WT^, *Col1a2*^OIM^ and previously analyzed E14.5 WT (E14.5^WT^) forelimbs (**Figure 6C**). Volcano plot analysis was used to visualize differences in ECM protein abundance between phenotypes (**Figure 6D**). Intrasample ratios for specific collagen isoforms were calculated using raw intensities (**Figure 6E**).

#### Brain and forelimb quantitative analysis

Similar to COL1A2 OIM forelimb quantitative analysis, LFQ intensities in the CS and IN fractions were normalized using **Eq. 2** and **Eq. 4**. Normalized LFQ intensities were used for volcano plot analysis (**Figure 7D**) and Pearson correlation coefficients (**Figure 7E**) to identify matrisome differences in developing WT brain and forelimb tissues. Intrasample ratios for specific collagen isoforms and other ECM proteins were calculated using raw intensities (**Figure 7F**). GO analysis of E14.5 forelimb and brain tissue (**Figure 7G**) was conducted using g:Profiler. ECM proteins that were more abundant or exclusive in either the brain or forelimb tissue were analyzed separately, and specific GO terms were reported along with corresponding *p*-values.

---

**Figure S1.**
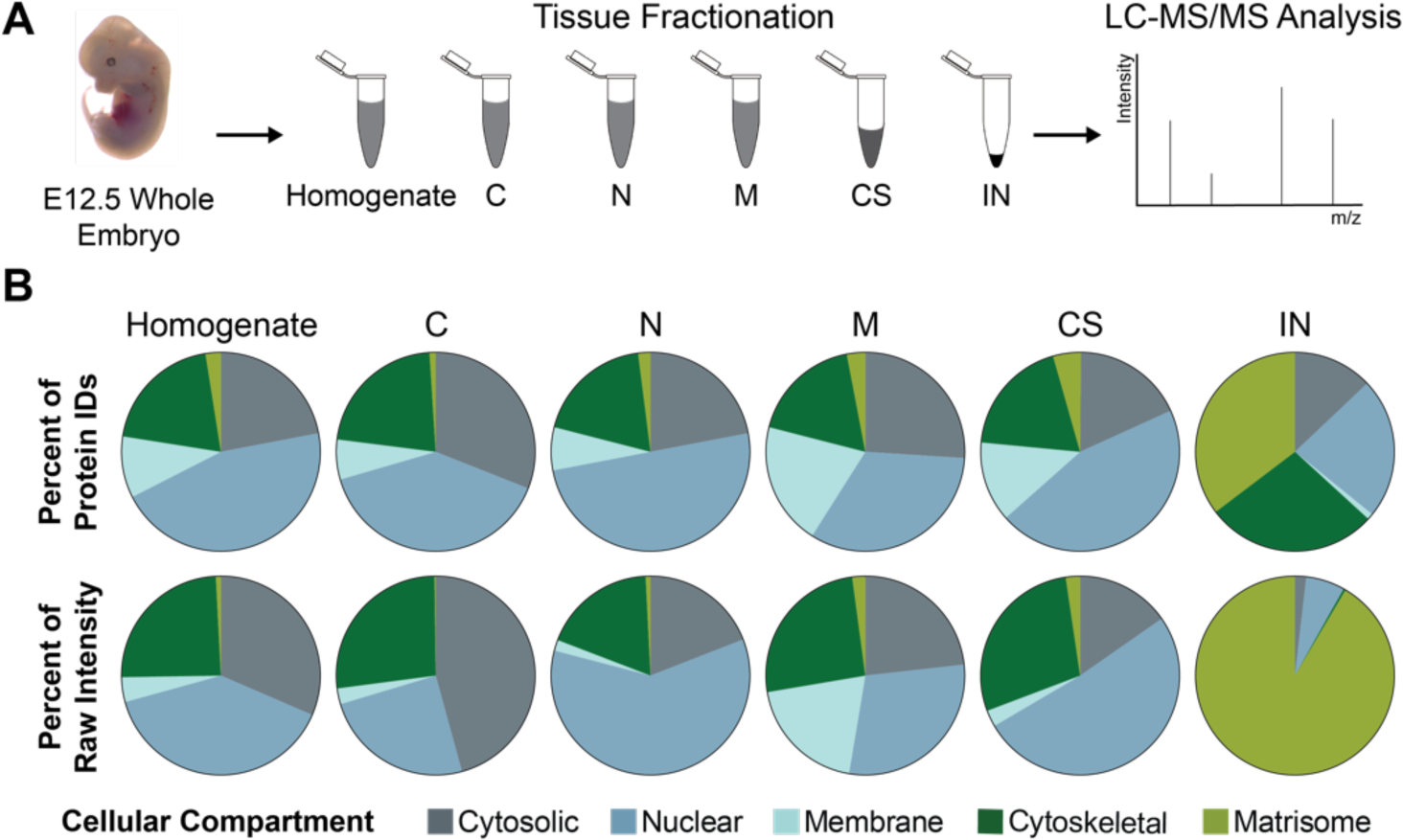
Proteomic analysis of fractionated E12.5 whole murine embryos. **(A)** Experimental workflow combining tissue fractionation with LC-MS/MS to investigate the matrisome coverage in each fraction. **(B)** The distribution of protein IDs and raw protein intensities, categorized by cellular compartment as defined by (Saleh, et al., 2019a), in homogenate, cytosolic (C), nuclear (N), membrane (M), cytoskeletal (CS) and insoluble (IN) fractions. There was minimal matrisome identification prior to tissue fractionation (average, *n*=2 biological replicates).

**Figure S2.**
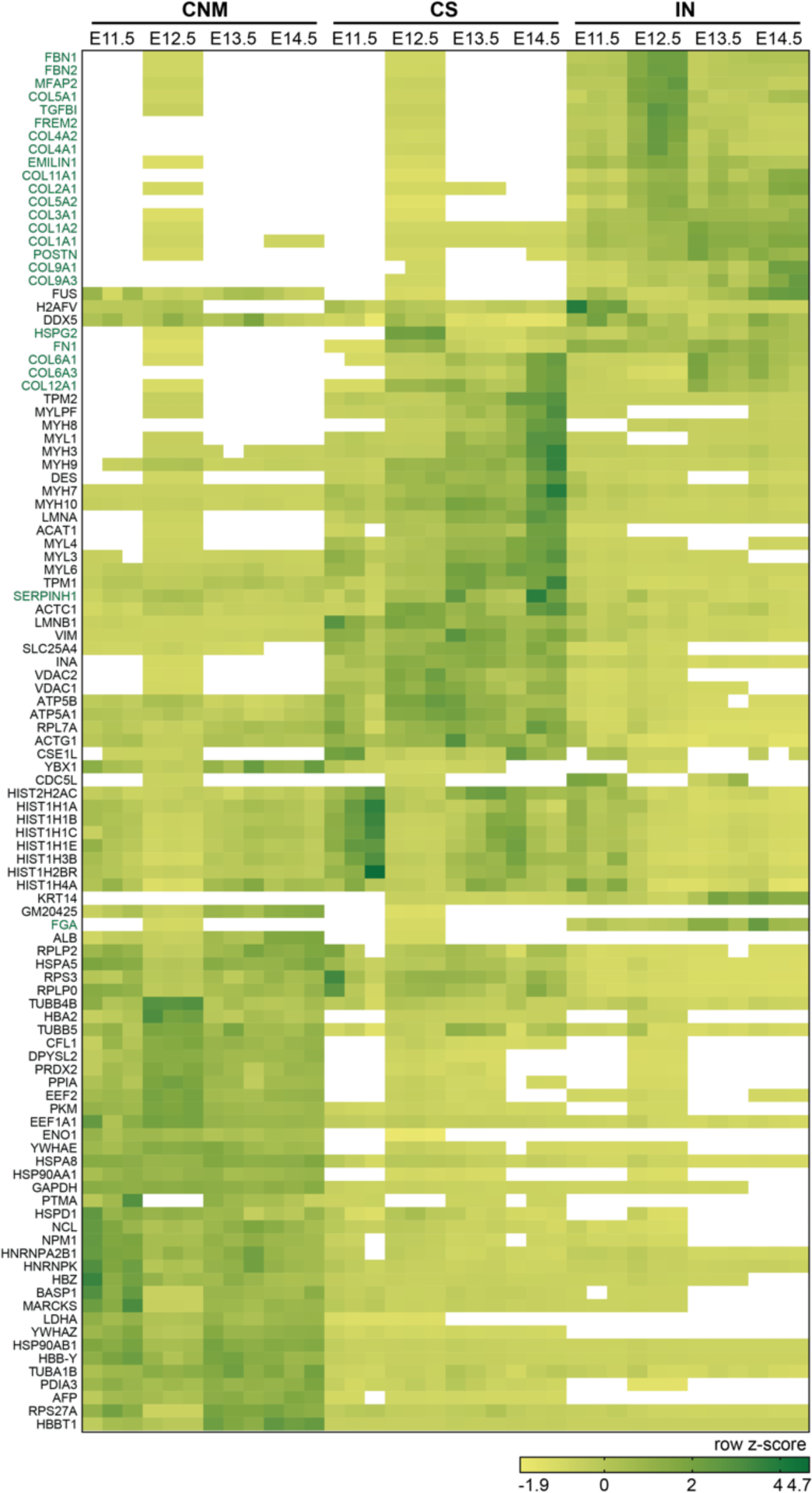
Unbiased hierarchal clustering of proteins identified in fractionated murine embryos. The 50 most abundant proteins in the cytosolic/nuclear/membrane (CNM), cytoskeletal (CS) and insoluble (IN) fractions of E14.5 embryos are shown. The row z-score was calculated across all timepoints and fractions for each protein. Unbiased clustering analysis, based on Pearson Correlation coefficients, showed that the majority of matrisome proteins (green text) were identified in the IN fraction. White boxes signify zero intensity values. Each column represents a biological replicate, with *n*=3 replicates per timepoint.

**Figure S3.**
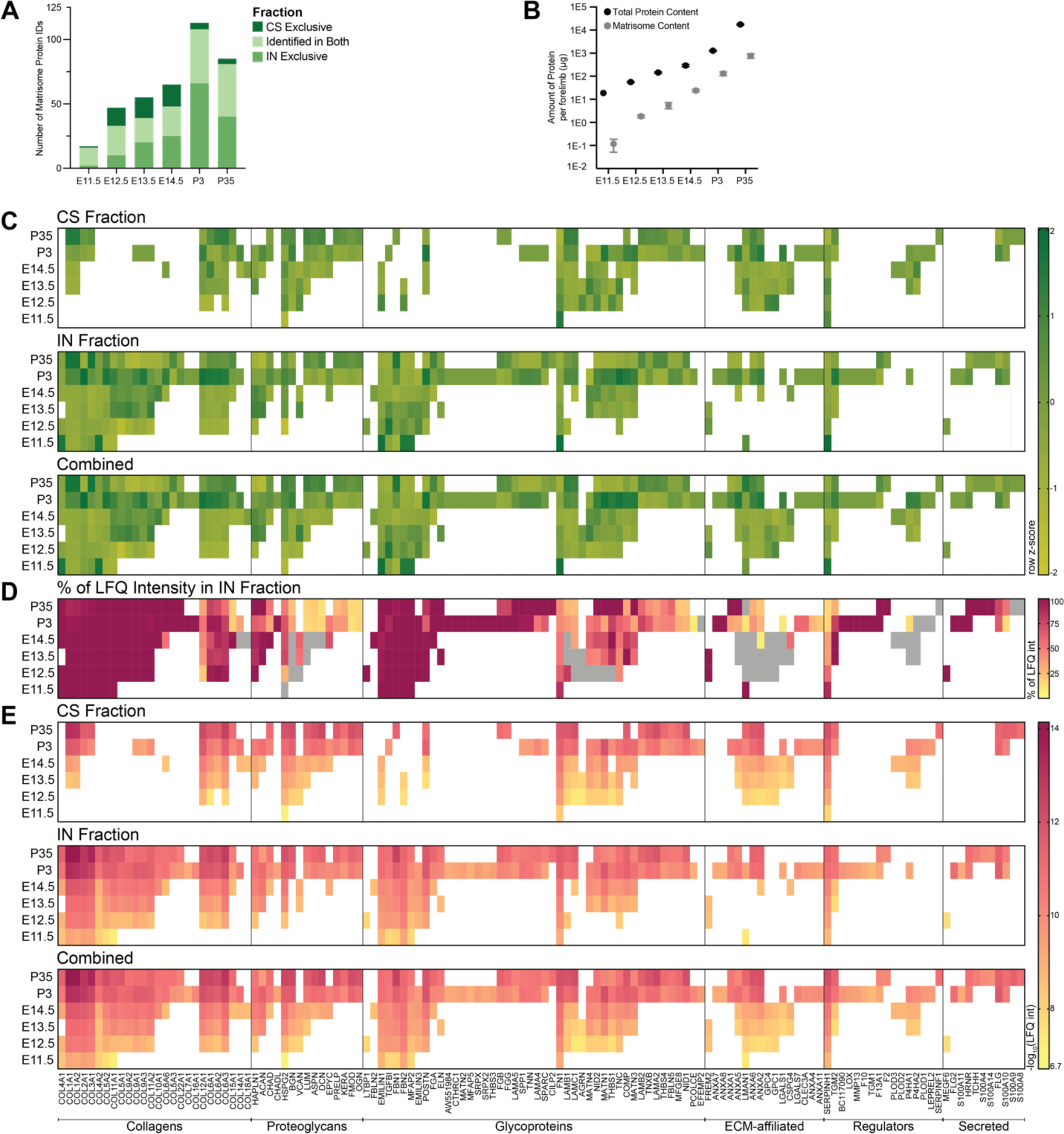
Comparative analysis of matrisome dynamics between fraction specific and combined LC-MS/MS LFQ intensities. **(A)** The number of ECM proteins identified exclusively in CS or IN fractions or distributed across both. **(B)** The average amount of total protein (black) and matrisome (grey) in one forelimb. Two-way ANOVA revealed the amount of total protein and matrisome content significantly increased as a function of development (*p*<0.0001). (**C**) Row z-scores were calculated for the normalized LFQ intensities in the CS (top) and IN (middle) fractions, separately, and clustered based on matrisome classification. By combining CS and IN intensities (bottom), the number of ECM protein identifications increased, but overall matrisome dynamics did not change. (**D**) The percentage of combined LFQ intensity (**C**, bottom) from the IN fraction (**C**, middle) indicates that there are differences in the amount of each ECM protein extracted in the CS fraction. Percentages were plotted as the average across biological replicates. Grey boxes denote protein intensities identified exclusively in the CS fraction (**C**, top). (**E**) LFQ intensities of ECM proteins quantified in CS and IN fractions were normalized individually (top and middle), then combined (bottom; see **Methods**). For heat map analysis, intensities were log10-transformed and plotted as the average of *n*=3 biological replicates. White boxes denote that protein was not identified at that timepoint.

**Figure S4.**
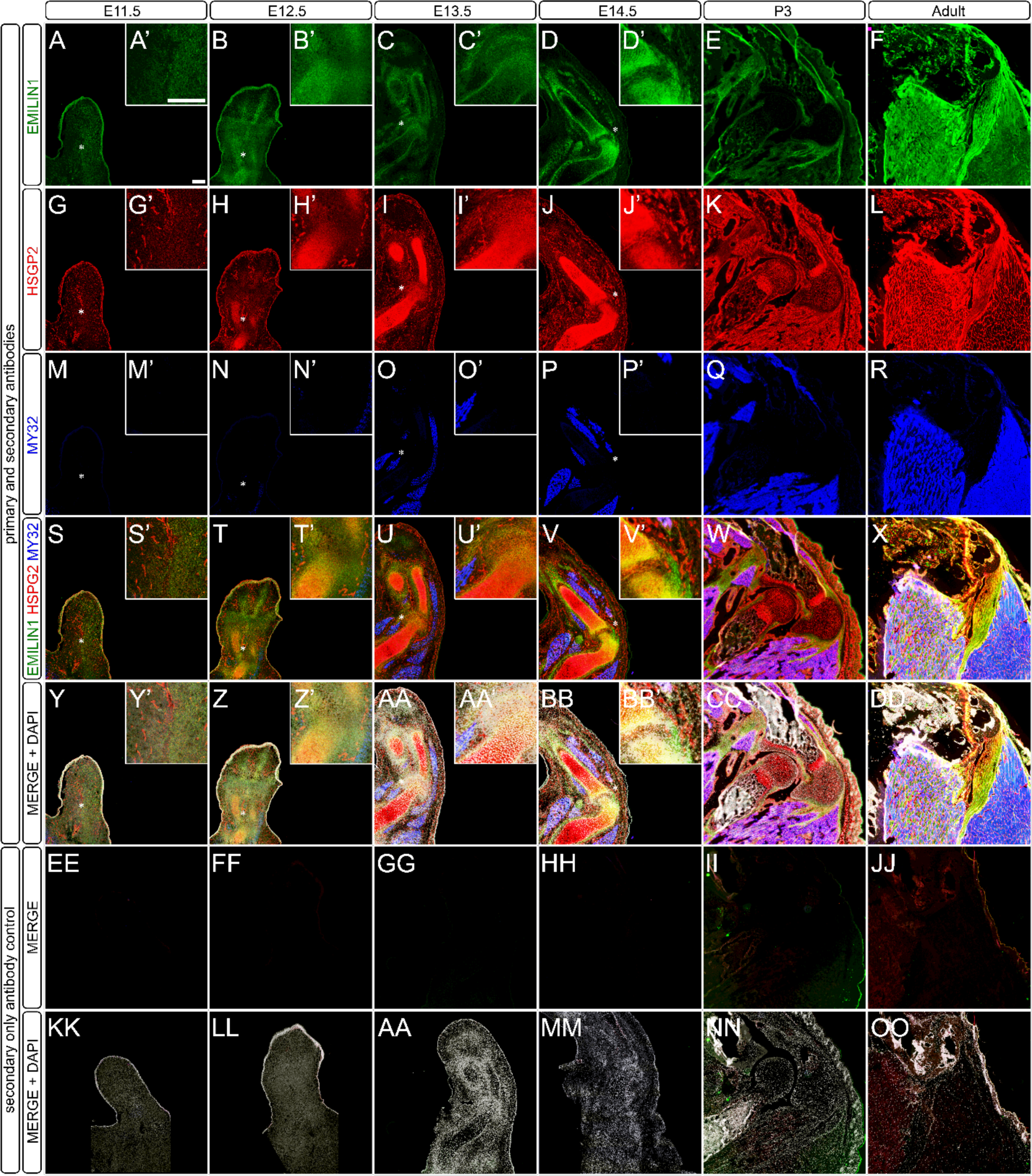
Spatiotemporal distribution of elastin microfibril interfacer 1 (EMILIN1) and perlecan (HSPG2) during forelimb development show differential patterning of proteins. **(A-DD)** Cryosections from E11.5-E14.5, P3 and adult were stained with antibodies against: **(A-F, A’-D’)** EMILIN1 (green); **(G-L, G’-J’)** perlecan (HSPG2; red); **(M-R, M’-P’)** myosin heavy chain, a marker for differentiated skeletal muscle (MY32; blue); **(S-X, S’-V’)** merge (green, red and blue); and **(Y-AD, Y’-AB’)** merge with DAPI (grey). **(EE-OO)** Secondary antibody only negative controls: **(EE-JJ)** merge; and **(KK-OO)** merge with DAPI. Insets (indicated with’) are a 3× enlargement of the forelimb containing the nascent elbow for E11.5-E14.5 at the location indicated with *. Scale bars = 200 µm.

**Figure S5.**
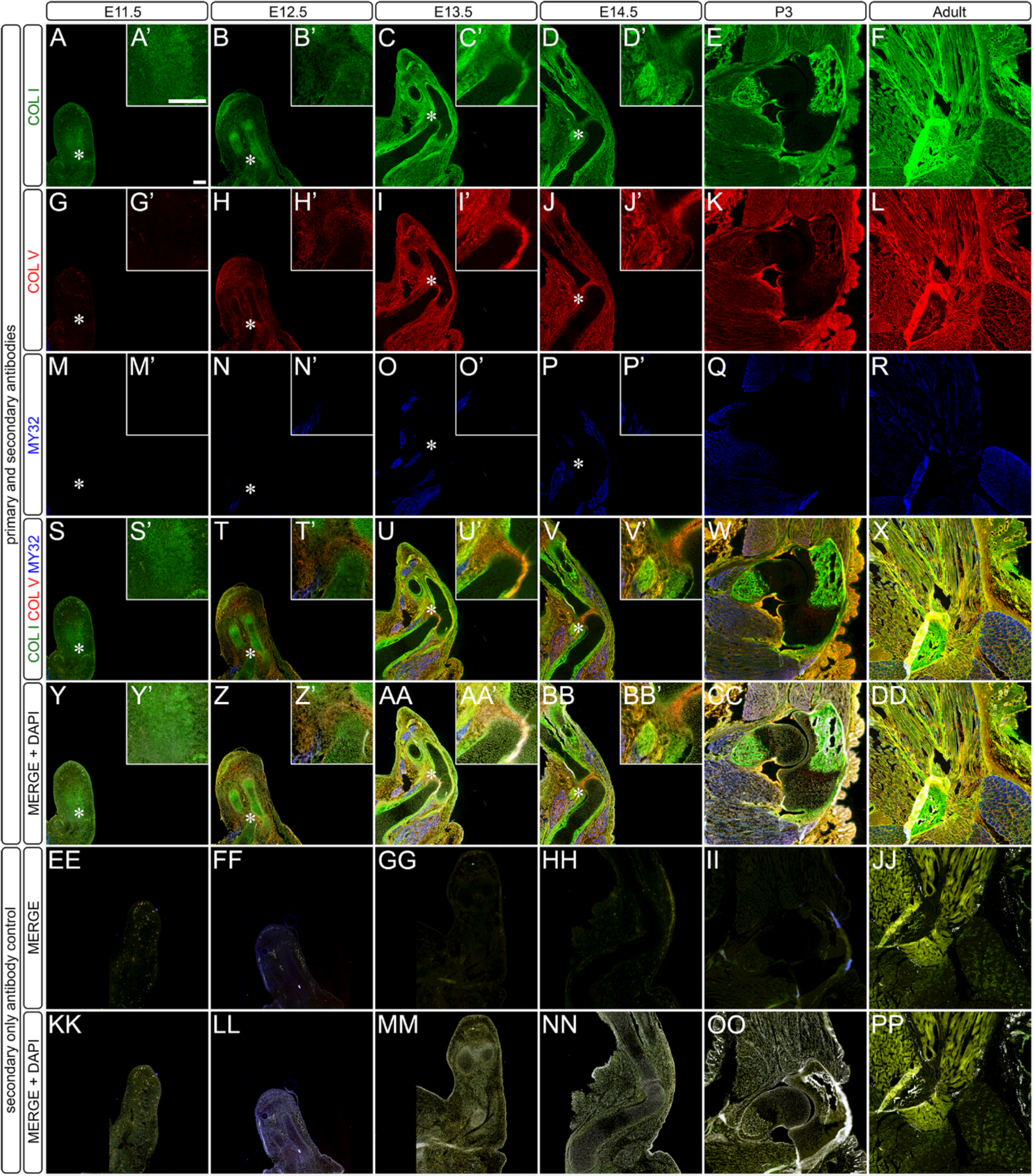
Spatiotemporal distribution of collagens, type I and V, during forelimb development show differential patterning of proteins. **(A-BB)** Cryosections from E11.5 – E14.5, P3 and adult were stained with antibodies against: **(A-F, A’-D’)** type I collagen (COL I; green); **(G-L, G’-J’)** type V collagen (COL V; red); **(M-R, M’-P’)** myosin heavy chain, a marker for differentiated skeletal muscle (MY32; blue); **(S-X, S’-V’)** merge (green, red, blue); and **(Y-DD, Y’-BB’)** merge with DAPI (grey). **(EE-OO)** Secondary antibody only negative controls: **(EE-JJ)** merge; and **(KK-OO)** merge with DAPI. Insets (indicated with’) are a 3× enlargement of the region containing the nascent elbow (*) for E11.5-E14.5. Scale bars=200 µm.

**Figure S6.**
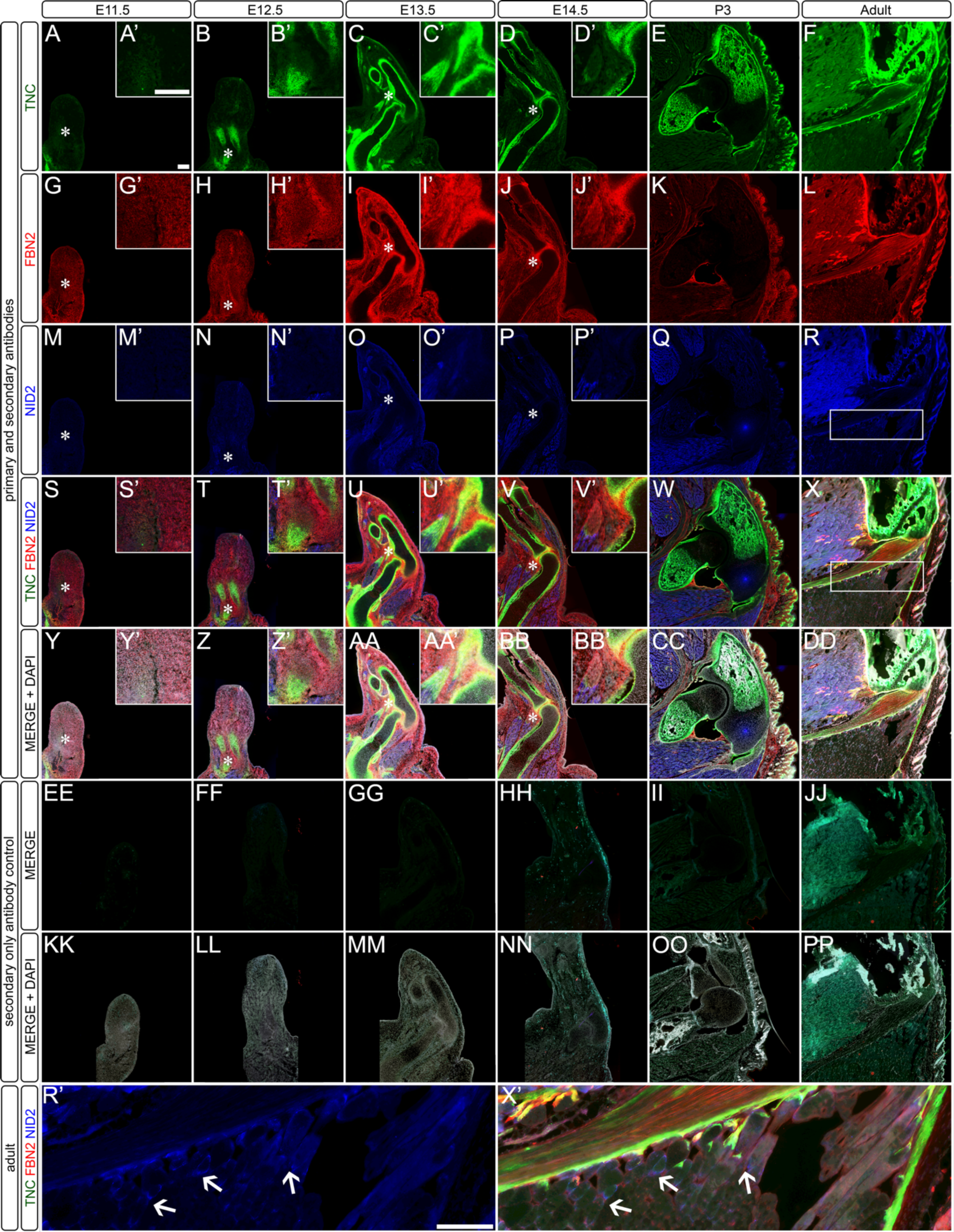
Spatiotemporal distribution of tenascin-C, fibrillin-2 and nidogen-2 during forelimb development show differential patterning of proteins. **(A-BB)** Cryosections from E11.5 – E14.5, P3 and adult were stained with antibodies against: **(A-F, A’-D’)** tenascin-C (TNC; green); **(G-L, G’-J’)** fibrillin-2 (FBN2; red); **(M-R, M’-P’)** nidogen-2 (NID2; blue); **(S-X, S’-V’)** merge (green, red, blue); and **(Y-DD, Y’-BB’)** merge with DAPI (grey). **(EE-OO)** Secondary antibody only negative controls: **(EE-JJ)** merge; and **(KK-OO)** merge with DAPI. Insets (indicated with’) **(R’)** NID2 and **(X’)** TNC/FBN2/NID2 merge channels showed punctate staining (arrow) of NID2 in the adult. White box in **(R)** and **(X)** highlighted the inset location. Other insets are a 3× enlargement of the region containing the nascent elbow (*) for E11.5– E14.5. Scale bars=200 µm.

**Figure S7.**
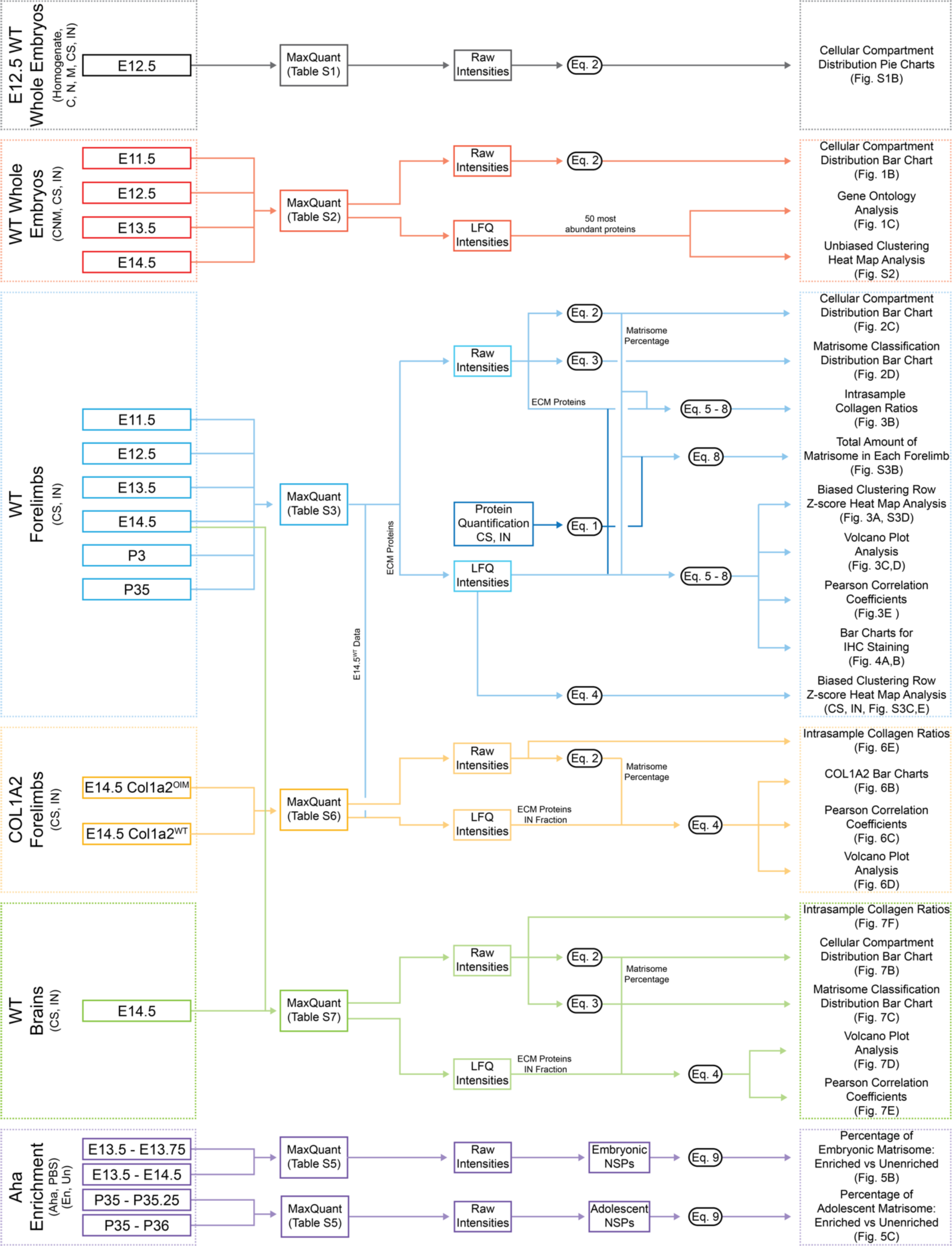
Workflow of data analysis.

